# Two interacting transcriptional coactivators cooperatively control plant immune responses

**DOI:** 10.1101/2021.03.21.436112

**Authors:** Huan Chen, Min Li, Guang Qi, Ming Zhao, Longyu Liu, Jingyi Zhang, Gongyou Chen, Daowen Wang, Fengquan Liu, Zheng Qing Fu

## Abstract

The phytohormone salicylic acid (SA) plays a pivotal role in plant defense against biotrophic and hemibiotrophic pathogens. Genetic studies have identified NPR1 and EDS1 as two central hubs in plant local and systemic immunity. However, it is unclear how NPR1 orchestrates gene regulation and whether EDS1 directly participates in transcriptional reprogramming. Here we show that NPR1 and EDS1 synergistically activate *Pathogenesis-Related* (*PR*) genes and plant defenses by forming a protein complex and co-opting with Mediator. In particular, we discover that EDS1 functions as an autonomous transcriptional coactivator with intrinsic transactivation domains and physically interacts with the CDK8 subunit of Mediator. Upon SA induction, EDS1 is directly recruited by NPR1 onto the *PR1* promoter via physical NPR1-EDS1 interactions, thereby potentiating *PR1* activation. We further demonstrate that EDS1 stabilizes NPR1 protein and NPR1 transcriptionally upregulates *EDS1* in plant-pathogen interactions. Our results reveal an elegant interplay of key coactivators with Mediator and elucidate novel molecular mechanisms for activating transcription during immune responses.

## Introduction

Plant-pathogen interactions have enabled plants to evolve a sophisticated and multifaceted immune system for defending against pathogen attacks^1^. Recognition of conserved pathogen-associated molecular patterns (PAMPs) by extracellular pattern recognition receptors in plants stimulates PAMP-triggered immunity (PTI). However, successful pathogens deploy a suite of virulence effectors to attenuate or dampen PTI, resulting in effector-triggered susceptibility. During host-pathogen coevolution, plants have developed resistance (R) proteins to specifically recognize pathogen-delivered effectors through direct interaction or indirect recognition by detecting the activities of pathogen effectors^2^, thus inducing a robust defense, termed effector-triggered immunity (ETI). Most R proteins belong to a large family of intracellular immune receptors known as nucleotide-binding (NB), leucine-rich repeat (LRR) receptor (NLR) proteins with a variable N terminal Drosophila Toll, mammalian interleukin-1 receptor (TIR)^3^ or coiled-coil (CC) domain^4^. Activation of PTI or ETI results in the generation of mobile signals that are transported from local infected tissue to distal uninfected parts^5^, inducing systemic acquired resistance (SAR), which is a long-lasting and broad-spectrum resistance against related or unrelated pathogens^6^.

The plant defense hormone salicylic acid (SA), as a small phenolic compound, plays a pivotal role in plant defense against biotrophic pathogens such as the oomycete pathogen *Hyaloperonospora arabidopsis* and hemibiotrophic pathogens such as the bacterial pathogen *Pseudomonas syringae*^7^. Pathogen-induced SA not only accumulates in infected local leaves but also in uninfected systemic tissues. As a consequence, SA is an essential signaling molecule for the activation of local defense and SAR^8-10^. Exogenous application of SA or its active analogues is sufficient to activate plant defense responses by inducing massive transcriptional reprograming to relocate energy for defense instead of growth^11,12^.

NONEXPRESSER OF PR GENES1 (NPR1) was identified through genetic screens for *Arabidopsis* mutants that cannot activate the expression of *PR* genes, which encode proteins with antimicrobial activities^13^. Similar to NPR3 and NPR4, NPR1 binds SA and functions as an SA receptor^14-17^. Before pathogen infection, NPR1 is sequestered in the cytosol as oligomers, which are crucial for protein homeostasis^18^. Upon pathogen challenge, oligomeric NPR1 is reduced into active monomers by SA-induced redox changes, and NPR1 monomers enter the nucleus^19^. As a transcriptional coactivator, NPR1 interacts with TGA and TCP transcription factors (TFs) and facilitates the expression of *PR* genes^20-22^. In addition to *PR* genes, NPR1 also controls the expression of the vast majority of other SA-responsive genes^23^. Therefore, it is believed that NPR1 functions as a master regulator of SA signaling.

ENHANCED DISEASE SUSCEPTIBILITY1 (EDS1) has been shown to be indispensable for TIR-NLR protein-dependent ETI, plant basal defense and SAR^24-28^. In addition to its association with numerous R proteins^26^, EDS1 physically interacts with PHYTOALEXIN DEFICIENT4 (PAD4) or SENESCENCE ASSOCIATED GENE101 (SAG101)^29^. Distinct EDS1-PAD4 and EDS1-SAG101 complexes are essential for different R protein-mediated ETI^30^. EDS1 and its partners have been shown to affect the expression of numerous pathogen-responsive genes^31^, but it remains unclear how EDS1 promotes downstream transcriptional reprogramming to trigger a series of immune responses.

In this study, we show that NPR1 and EDS1 interact with each other to form a protein complex and synergistically activate plant immunity via SA signaling. We demonstrate that EDS1 possesses transcriptional activation activity and serves as an acidic transcriptional coactivator, which is directly involved in transcriptional reprogramming by interacting with a component of the Mediator complex, cyclin-dependent kinase 8 (CDK8). Moreover, we find that upon SA induction, NPR1 directly recruits EDS1 to the *PR1* promoter to facilitate the expression of *PR1*. Furthermore, we identify a positive feedback loop, in which NPR1 directly upregulates *EDS1* transcription and EDS1 stabilizes NPR1 protein in plant-pathogen interactions. Our study revealed a unique mechanism, in which two interacting transcriptional coactivators co-opts with Mediator and cooperatively control transcriptional reprograming to activate plant defense responses.

## Results

### NPR1 physically interacts with EDS1 to form a protein complex

Both NPR1 and EDS1 function as central hubs in plant immunity^32,33^, and they have also been identified as targets of pathogen effectors^26,34^. In a yeast-two hybrid (Y2H) screen, we have identified EDS1 as an NPR1 interactor. NPR1 specifically interacted with EDS1, but not with PAD4 or SAG101, two other members of the EDS1 family of lipase-like proteins in Y2H assays (Fig. 1a and Supplementary Fig. 1a). The specific NPR1-EDS1 interaction was then confirmed by *in vitro* pull-down assays, where Thioredoxin (Trx)-His_6_-NPR1 bound glutathione S-transferase (GST)-EDS1, but not GST-PAD4 or GST (Fig. 1b). Their interaction *in planta* was determined using co-immunoprecipitation (Co-IP) assays in *Nicotiana benthamiana*, in which EDS1-Myc was co-immunoprecipitated with NPR1-FLAG (Fig. 1c). Using a bimolecular luminescence complementation (BiLC) assay in *N. benthamiana*, the *in planta* interaction between NPR1 and EDS1 was further confirmed (Fig. 1d). Taken together, these data demonstrate that NPR1 interacts with EDS1 *in vitro* and *in vivo*.

**Fig. 1.**
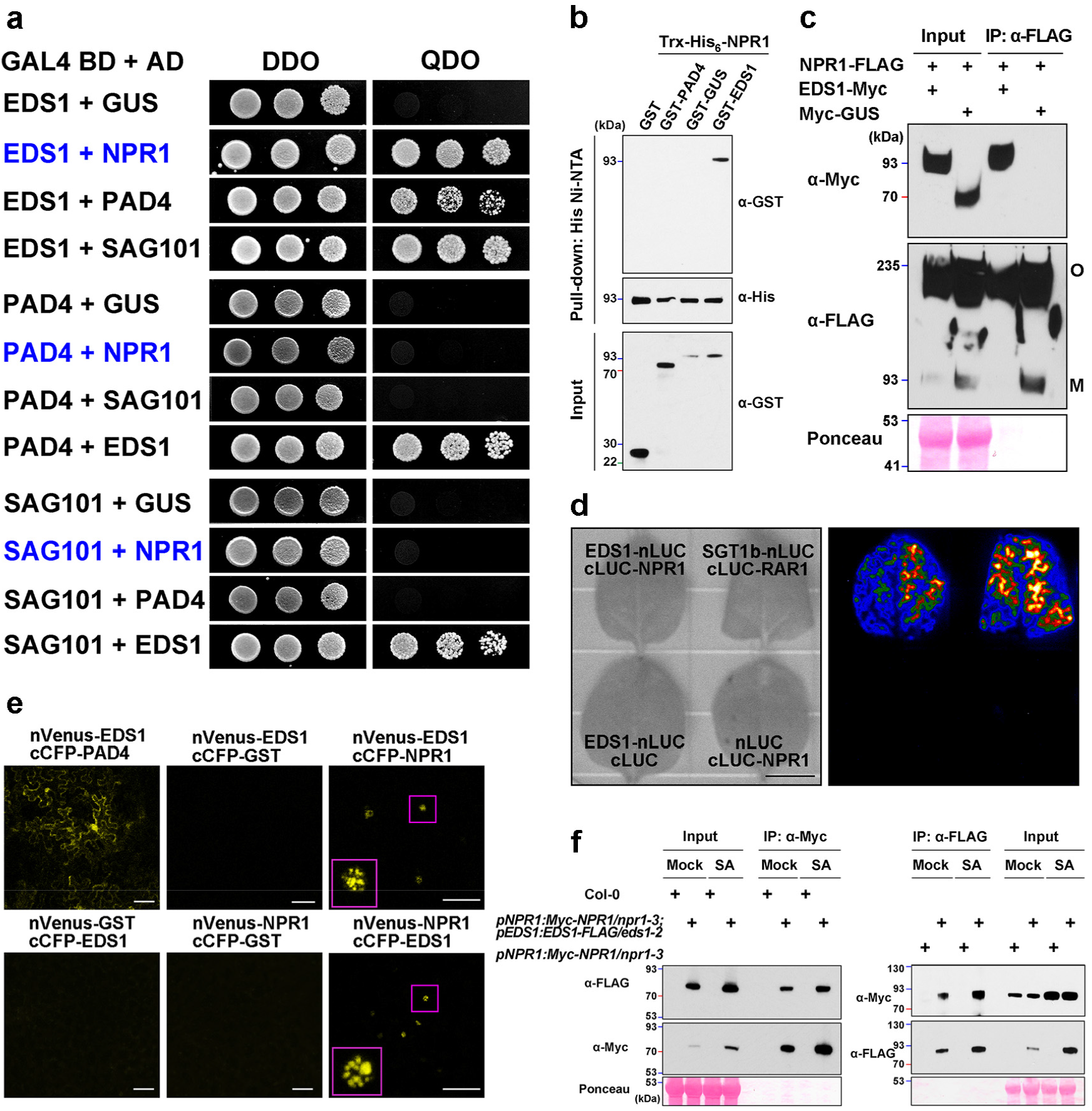
NPR1 directly interacts with EDS1. **a**, NPR1 interacts with EDS1 in Y2H assays. The growth of yeast strains on nonselective double dropout medium (DDO) and selective quadruple dropout medium (QDO) is shown. GAL4 BD, GAL4 DNA-binding domain; AD, activation domain. **b**, NPR1 interacts with EDS1 in the *in vitro* pull-down assays. Trx-His_6_-NPR1 was used to pull down GST and GST fusion proteins. Trx-His_6_-NPR1 and GST fusion proteins were detected by western blotting with anti-His and anti-GST antibodies, respectively. **c**, NPR1 interacts with EDS1 in *N. benthamiana*. The *NPR1-3FLAG* under the control of its native promoter was transiently expressed with *EDS1-Myc* under the control of its native promoter or *Myc-GUS* under the control of the 35S promoter in *N. benthamiana*. Co-IP assay was performed using anti-FLAG magnetic beads. O, oligomeric NPR1; M, monomeric NPR1. **d**, NPR1 interacts with EDS1 in BiLC assays. The indicated vectors were coexpressed in *N. benthamiana* leaves and luciferase complementation imaging assays were performed. Scale bar, 1 cm. **e**, NPR1 interacts with EDS1 in nuclei in BiFC assays. *N. benthamiana* was co-transformed with indicated constructs. Magnified nuclear body is shown in red box. Scale bars, 150 µm. **f**, NPR1 interacts with EDS1 in *Arabidopsis*. The two-week-old *Arabidopsis* seedlings were treated with 0.5 mM SA or water (Mock) for 9 h, and total protein extract was subject to Co-IP assays using Myc-Trap_MA or anti-FLAG magnetic beads. Ponceau S staining of RuBisCo is used for confirmation of equal loading. Protein sizes marked on the left are in kDa.

We next carried out a bimolecular fluorescence complementation (BiFC) assay to check the subcellular localization of NPR1-EDS1 complex by transiently expressing these two proteins in *N. benthamiana* using agroinfiltration. Compared with the EDS1-PAD4 complex that was detected mainly in the nucleus and cytoplasm, the observed NPR1 association with EDS1 in the nucleus apparently formed nuclear bodies (Fig. 1e), which most likely act as the sites for accelerating gene activation or repression^35^. These data imply that the primary function of the NPR1-EDS1 protein complex is to regulate the expression of plant defense genes. In order to validate the native NPR1-EDS1 interaction in *Arabidopsis*, we produced transgenic lines expressing the *EDS1* native promoter-driven EDS1-FLAG in Col-0 *eds1-2* mutant (p*EDS1*:*EDS1-FLAG*/*eds1-2*) and crossed it with p*NPR1*:*Myc-NPR1*/*npr1-3* transgenic lines to obtain the p*NPR1*:*Myc-NPR1*/*npr1-3*; p*EDS1*:*EDS1-FLAG*/*eds1-2* plants. In reciprocal Co-IP experiments, we detected that SA enhances the NPR1-EDS1 association possibly due to the increased protein levels of NPR1 and EDS1 after SA treatment (Fig. 1f). Altogether, these data suggest that SA induces the accumulation of NPR1-EDS1 protein complex within the nuclear bodies to facilitate the expression of plant defense genes.

We additionally conducted Y2H assays to identify the domain of EDS1 that is necessary for its interaction with NPR1. Several EDS1 fragments including the EP (EDS1 and PAD4-defined) domain, the Lipase-like domain, the helical region encompassing amino acid residues from 310 to 350 (310-350), and a coiled-coil domain (358-383) (Supplementary Fig. 1b) were tested based on the secondary and crystal structures^25,29^. The results showed that the helical region (310-350) is sufficient and necessary for EDS1 to interact with NPR1. To narrow down the interacting region in the helical structure, we divided it into two alpha helices (310-330 and 331-350). EDS1 lacking residues 310 to 330 (Δ310-330) failed to interact with NPR1, while the minimal region (310-330) exhibited obvious interaction. Therefore, the minimal alpha helix (310-330) in EDS1 is necessary and sufficient for the interaction with NPR1. Based on the crystal structure of EDS1^29^, this minimal alpha helix (310-330) is located on the surface of the N-terminal domain of EDS1 (Supplementary Fig. 1c), further supporting the critical role of this region for EDS1-NPR1 interaction.

Conversely, we also generated different truncations of NPR1 and identified several domains of NPR1 that are involved in NPR1-EDS1 interaction (Supplementary Fig. 1d,e). Intriguingly, we found that the BTB/POZ (for Broad complex, Tramtrack, and Bric-a-brac/Pox virus and Zinc finger) domain, the ankyrin repeats (ANK) and an important C-terminal domain (CTD) interact with EDS1 in Y2H assays. BTB/POZ and ANK motifs are well known as protein-protein interaction motifs in a number of proteins in mammals and plants^13,36,37^. To further decipher whether the interaction of NPR1 with EDS1 is relevant to NPR1’s function, we investigated the interaction of EDS1 with mutant npr1 protein encoded by several *npr1* alleles (i.e., *npr1-2, nim1-2, npr1-1*, and *npr1-5*) that are compromised in SA signaling and SAR induction^38-41^. Consistently, *npr1-2* (C150Y) mutation in BTB domain or other point mutations in ANK region such as *nim1-2* (H300Y), *npr1-1* (H334Y) and *npr1-5* (P342S) completely lost the ability to interact with EDS1 (Supplementary Fig. 1d), indicating that the interaction of EDS1 with NPR1 is important for the function of NPR1. In addition, it has been revealed that the CTD overlapping a repression region of NPR1^22^ is probably involved in SA perception^16,42^. Collectively, these findings suggest that multiple regions of NPR1 are required for the dynamic interaction with EDS1 in plant immune responses.

### NPR1 and EDS1 cooperatively activate plant immunity

EDS1 plays an essential role in ETI triggered by TIR-NLR proteins^24^. EDS1 is required for recognition of the *Pseudomonas syringae* pv. tomato avirulence effector Rps4 (*Pst* AvrRps4) by the nuclear R protein pair RESISTANT TO RALSTONIA SOLANACEARUM1S (RRS1S)-RESISTANT TO PSEUDOMONAS SYRINGAE (RPS4)^43^. To dissect the roles of the NPR1-EDS1 interaction in controlling plant immunity, genetic interactions were analyzed between recessive *npr1* mutant alleles (*npr1-2* and *npr1-3*)^13^ and the null *eds1-2* allele^31^ in the *Arabidopsis thaliana* ecotype Col-0 background. In comparison to *eds1-2*, the *npr1-2* and *npr1-3* are moderately susceptible to *Pst* DC3000 *avrRps4*, whereas two homozygous transgenic lines overexpressing N-terminal green-fluorescent protein (GFP) tagged NPR1 (35S:*GFP-NPR1* #11 and #36) were robustly resistant to the avirulent pathogen (Fig. 2a). This suggests that NPR1 prominently contributes to RRS1S/RPS4-mediated ETI. To further test the function of nuclear NPR1 in ETI, we examined the resistance of the transgenic plants expressing *NPR1-GFP* and its nuclear localization signal (NLS) mutant form *NPR1 (nls)-GFP* to *Pst* DC3000 *avrRps4*. As shown in Supplementary Fig. 2a, the enhanced ETI conferred by *NPR1-GFP* was completely lost in *NPR1 (nls)-GFP* transgenic plants, revealing that nuclear NPR1 contributes to ETI likely through transcriptional regulation.

**Fig. 2.**
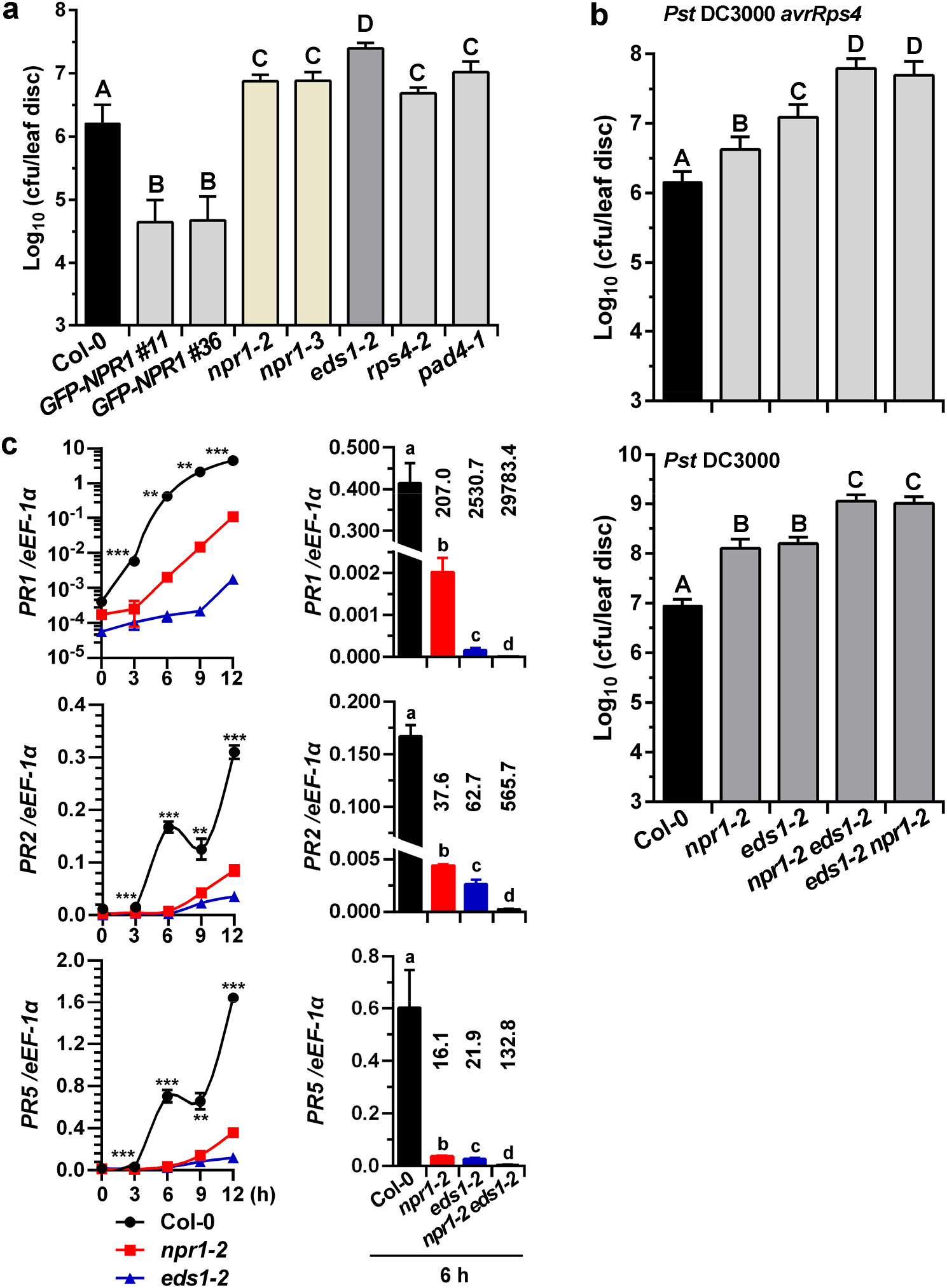
Genetic and molecular interactions of NPR1 with EDS1. **a**, NPR1 contributes to ETI. Growth of *Pst* DC3000 *avrRps4* on Col-0, different mutants and transgenic *Arabidopsis* overexpressing *GFP-NPR1* under the control of the *CaMV* 35S promoter. **b**, NPR1 and EDS1 additively activate immune responses. Leaves from soil-grown *Arabidopsis* (**a**,**b**) were hand-infiltrated with indicated bacterial suspensions (OD_600_ = 0.0005) and bacterial titers were measured at 2 d post-inoculation (dpi). CFU, colony-forming units. **c**, NPR1 and EDS1 synergistically upregulates *PR* genes. Leaves from 4-week-old plants inoculated with *Pst* DC3000 *avrRps4* (OD_600_ = 0.01) were collected at indicated time points and *PR* gene expression was checked using real-time qPCR. Expression of *PR1* was plotted on a log_10_ scale; gene expression levels were normalized against the constitutively expressed *eEF-1α*. Right panel, the expression of *PR* genes at 6 h after pathogen infection. Notably, the numbers above the error bars indicate the fold change of gene expression compared with Col-0. Error bars represent standard deviation (SD). n = 6 biologically independent samples (**a**,**b**); n = 4 biologically independent samples (**c**). Statistically significant differences are indicated by different lowercase letters (ANOVA, *P* < 0.05) or shown between Col-0 and single mutant (*npr1-2* or *eds1-2*) plants (*t*-test, **, *P* < 0.01; ***, *P* < 0.001).

In further epistasis analysis, the double mutants (*npr1-2 eds1-2* and *eds1-2 npr1-2*) obtained from two reciprocal crosses (*npr1-2* × *eds1-2* and *eds1-2* × *npr1-2*) were more susceptible to *Pst* DC3000 *avrRps4* and *Pst* DC3000 than either *npr1-2* or *eds1-2* single mutants (Fig. 2b), demonstrating that NPR1 and EDS1 additively contribute to ETI and basal resistance. To determine whether EDS1 is involved in NPR1-mediated defense pathways, *NPR1-GFP*/*npr1-2* transgenic plants were crossed with *eds1-2* mutants and homozygous *NPR1-GFP*/*npr1-2*;*eds1-2* plants were identified and analyzed. *NPR1-GFP*/*npr1-2*;*eds1-2* plants exhibit a susceptibility somewhat less than that conferred by *eds1-2* (Supplementary Fig. 2b), suggesting that EDS1 functions both dependently and independently of NPR1 to regulate ETI. To further confirm the function of NPR1-EDS1 interaction in plant defense, we examined the susceptibility of these mutants to another avirulent pathogen *P. syringae* pv. *maculicola* (*Psm*) ES4326 *avrRpt2*, which activates ETI mediated by the CC-NLR protein RPS2 (ref.^44^). We found that the growth of *Psm* ES4326 *avrRpt2* in *eds1-2* is not significantly higher than wild-type plants. However, the pathogen growth in *npr1-2 eds1-2* or *eds1-2 npr1-2* was higher than either *npr1-2* or *eds1-2* (Supplementary Fig. 2c), indicating the cooperative contributions of NPR1 and EDS1 in RPS2-mediated ETI. Thus, these genetic interaction data are consistent with the hypothesis that NPR1 and EDS1 function as partners in diverse immune responses.

In addition to genetic interactions, the molecular function of the NPR1-EDS1 interaction was investigated. We used real-time quantitative PCR (qPCR) to monitor the time course expression of *PR* genes (*PR1, PR2*, and *PR5*), a subset of EDS1-induced *WRKY* genes and two EDS1-repressed genes (*DND1* and *ERECTA*) in ETI^31,45^. These *PR* genes and EDS1 target genes were mis-regulated in *npr1-2* and *eds1-2* in similar manners (Fig. 2c and Supplementary Fig. 2d). Interestingly, loss of *EDS1* function has a stronger effect than loss of *NPR1* function on expression of *PR* genes after pathogen infection (Fig. 2c), consistent with our bacterial growth data (Fig. 2a and Supplementary Fig. 2b). Importantly, the reduction in the expression of *PR* genes in the *npr1-2 eds1-2* was more pronounced than single mutants (Fig. 2c, right). These results indicate that the synergistic regulation of defense gene expression by NPR1 and EDS1 is essential for immune responses.

### NPR1 and EDS1 synergistically promote plant defenses via SA signaling

EDS1 can function upstream of SA in plant immunity because EDS1 contributes to pathogen-induced SA accumulation^46^. PR proteins have been considered as hallmarks of SA signaling^38,47,48^. Notably, the *eds1-2* mutation strongly inhibited *Pst* DC3000 *avrRps4*-induced *PR* gene expression (Fig. 2c); thus, EDS1 may also function downstream of SA. To study potential roles of EDS1 in SA signaling, the induction of *PR* genes in response to exogenous SA was examined in two *eds1* mutant alleles. The result from qPCR analysis showed that the expression of SA-induced *PR* genes was significantly decreased in the *eds1-2* rosette leaves from soil-grown plants and seedlings (Supplementary Figs. 3a,b). Moreover, SA-induced PR1 protein accumulation in Ws-0 *eds1-1* or Col-0 *eds1-2* was significantly attenuated and delayed (Fig. 3a), compared to the corresponding wild-type plants. Concomitantly, the gradually increased EDS1 protein over time was highly elevated by SA (Fig. 3a). Hence, the SA-induced EDS1 indeed plays a positive role in SA signaling to activate defense genes. We also examined SA-induced pathogen resistance in *eds1* mutants. In contrast to *npr1-2* plants, exogenous application of SA significantly rendered *eds1-1* and *eds1-2* plants resistant to *Pst* DC3000 (Fig. 3b), consistent with the idea that EDS1 functions upstream of SA. However, the SA-induced pathogen resistance in *eds1-1* or *eds1-2* was not as strong as that in wild-type plants (Fig. 3b). These findings are in agreement with our conclusion that EDS1 can function as a positive regulator of SA signaling in immune responses.

**Fig. 3.**
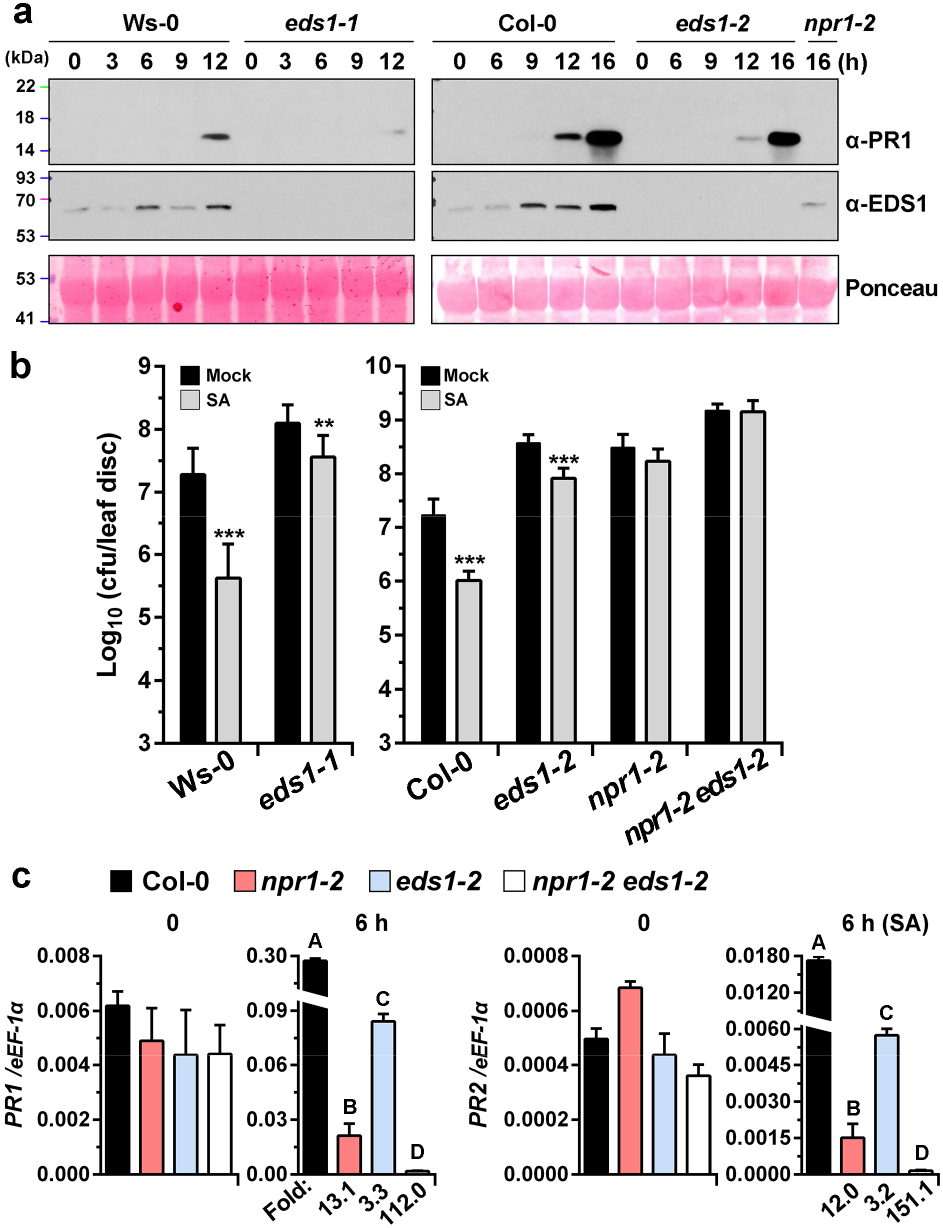
NPR1 and EDS1 function downstream of SA. **a**, EDS1 is a positive regulator of SA signaling. Total protein was prepared from leaf tissues of 4-week-old plants infiltrated with 0.3 mM SA and subjected to immunoblotting with indicated antibodies. This experiment is representative of at least two independent replicates. **b**, EDS1 contributes to SA-induced pathogen resistance. Plants were treated with soil drenches plus foliar sprays of 0.5 mM SA or water (Mock). After 24 h, leaves were inoculated with *Pst* DC3000 (OD_600_= 0.0005) and the *in planta* bacterial titers were determined at 3 dpi. Error bars represent standard error (SE); n = 3 biologically independent experiments. Statistical differences from Mock in each genotype are shown (*t*-test, **, *P* < 0.01; ***, *P* < 0.001). **c**, NPR1 and EDS1 synergistically upregulate *PR1* and *PR2*. Two-week-old seedlings grown on 1/2 MS media were exogenously treated with hydroponic 0.5 mM SA solution for 6 h. Total RNA was extracted and subjected to qRT-PCR. Error bars indicate SD; n = 4 biologically independent samples. Different letters indicate statistical differences (*P* < 0.01). Folds on the x-axis indicate fold reduction of gene expression compared with the value obtained in Col-0.

To investigate the functions of the NPR1-EDS1 interaction in SA signaling, we examined the expression of other SA-responsive and NPR1 target genes in *eds1-2* mutant. In addition to NPR1-dependent *PR* genes (Fig. 3a and Supplementary Fig. 3a,b), some *WRKY* genes, as well as several genes involved in pathogen-induced SA accumulation were significantly reduced in *eds1-2* compared with Col-0 after SA treatment (Supplementary Fig. 3c), suggesting that EDS1 and NPR1 upregulate the expression of a common set of SA-induced genes. To determine the contribution of EDS1 to the *PR* gene expression in the absence of NPR1, the expression of both *PR1* and *PR2* was examined in *npr1-2 eds1-2* in response to SA. Remarkably, the expression of *PR1* and *PR2* were dramatically reduced in *npr1-2 eds1-2* seedlings (Fig. 3c), compared to *npr1-2* or *eds1-2* seedlings. Strikingly, the fold change for the reduction of *PR1* expression in *npr1-2 eds1-2* (112.0-fold) was much greater than the product of the fold change in *npr1-2* (13.1-fold) and *eds1-2* (3.3-fold). Similar results were found with seedlings that were grown on media containing low concentrations of SA for a long-term treatment (Supplementary Fig. 3d). Therefore, these results demonstrate that NPR1-EDS1 interaction synergistically activates SA-mediated defense and pathogen resistance.

### The NPR1-EDS1 complex associates with specific chromatin regions upon SA induction

To further explore the effects of NPR1-EDS1 complex on the expression of defense genes, we conducted a series of chromatin immunoprecipitation (ChIP) assays. Multiple NPR1-interacting TGA factors^49-51^ and numerous WRKY TFs, that bind specifically to the W-box motif (TTGACC/T), have been shown to play essential roles in plant defense^52-54^. Based on these studies, we chose a set of promoter fragments that contain the common TGA motif (TGACG), the preferred TGA2-binding motif (TGACTT)^16^, or the W-box in our ChIP assays. As shown in Fig. 4a, Myc-NPR1 specifically associates with chromatin fragments at the *PR1* promoter in p*NPR1*:*Myc-NPR1*/*npr1-3* transgenic plants after SA treatment. In contrast, Myc-NPR1 did not significantly associate with the *PR1* promoter in the absence of SA, probably owing to constitutive protein degradation^55^ and the persistent existence of cytosolic oligomers under noninducing conditions^18^. Surprisingly, EDS1-FLAG in p*EDS1*:*EDS1-FLAG*/*eds1-2* transgenic lines and Myc-NPR1 bind almost identical sites on the *PR1* promoter (Fig. 4a). Moreover, NPR1 and EDS1 associated with the promoters of *PR2* and *PBS3* with similar enrichment profiles especially after SA treatment (Supplementary Fig. 4a,b). Together, these results demonstrate that EDS1 is a chromatin-associated protein and the NPR1-EDS1 complex associates with specific chromatin regions upon SA induction.

**Fig. 4.**
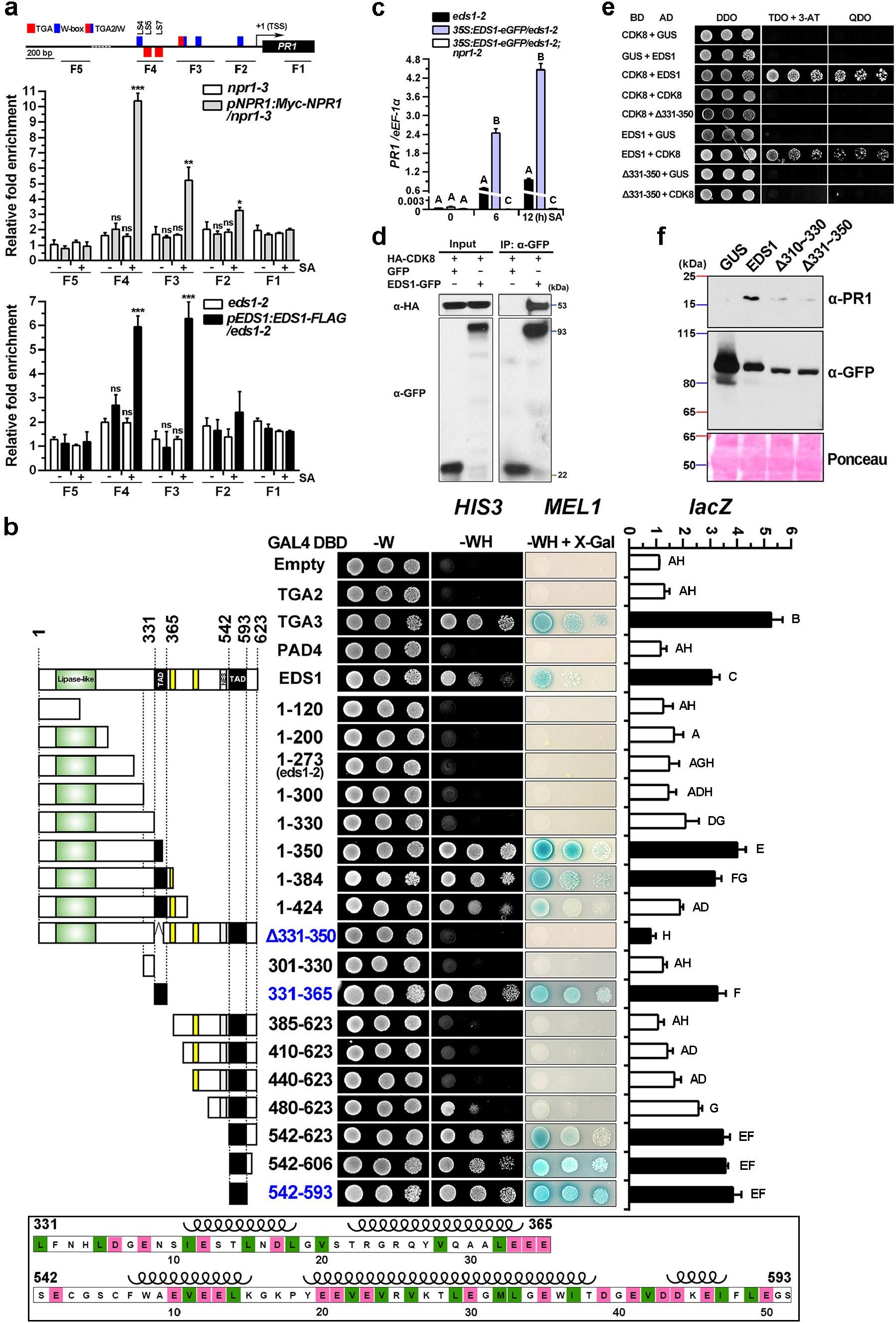
EDS1 functions as an acidic transcriptional coactivator. **a**, The association of NPR1 and EDS1 with the *PR1* promoter depends on SA. ChIP-qPCR analysis of NPR1 and EDS1 enrichment at *PR1* genomic loci was performed with anti-Myc or anti-FLAG antibody. Plants were harvested after 0.5 mM SA (+) or water (-) treatment for 9 h. Error bars indicate SD. n = 3 biologically independent samples. Significances of differences from the *npr1-2* treated with water are shown (*t*-test, *, *P* < 0.05; **, *P* < 0.01; ***, *P* < 0.001). Top panel, schematic representation of the *cis*-elements and chromatin fragments of amplicon in the *PR1* genomic region. Detailed positions of primers are described in Supplementary Table 1. LS4, equivalent to W-box (blue); LS5 and LS7, TGA motif (red); TGA2/W (chimeric color), TGA2 binding sites overlapping W-box; TSS, transcriptional start site. The TGA motifs with inverted consensus sequences are shown below. **b**, EDS1 functions as transcription activator with two autonomous TADs. Left panel, schematic illustration of EDS1 and the deletion mutants. Lipase-like domain (green); nuclear localization signal (yellow); NES, nuclear export signal (white); TAD, transactivation domain (solid). Middle panel, qualitative assay of yeast growth on selective media lacking tryptophan and histidine (-WH) and the media supplemented with X-α-Gal. Right panel, quantitative β-galactosidase analysis of LacZ activity. Bars denote SD of four biologically independent replicates (n = 4). Different letters indicate statistical differences (*P* < 0.05). Low panel, sequence of two TADs in EDS1. Acidic amino acid residues such as aspartic acid (Asp/D) and glutamic acid (Glu/E) are labeled in red. Hydrophobic residues including leucine (Leu/L), isoleucine (Ile/I), valine (Val/V) and methionine (Met/M) are shown in green. Coils refer to the positions of helices in the crystal structure. Numbers indicate amino acid positions. **c**, SA-induced EDS1 activates *PR1*. Expression of *PR1* was analyzed by qRT-PCR. Four-week-old soil-grown plants were infiltrated with 0.5 mM SA solution. Bars represent SD; n = 3 biologically independent samples. Different letters indicate significant differences (two-way ANOVA, *P* < 0.05). The statistical comparisons were made separately among different genotypes for each time point. **d**,**e**, EDS1 directly interacts with CDK8. *35S*:*HA-CDK8* was co-expressed with *35S*:*EDS1-GFP* or *35S*:*GFP* in *N. benthamiana*. Yeast cells were grown on DDO and selective triple dropout medium (TDO, without Leu, Trp and His) plus 1 mM 3-aminotriazole (3-AT) and QDO medium. **f**, Induction of PR1 affected by EDS1 deletion mutants. *35S:GUS-eGFP, 35S:EDS1-eGFP, 35S:EDS1 (*Δ*310∼330)-eGFP, 35S:EDS1 (*Δ*331∼350)-eGFP* were transformed into *eds1-2* young leaves by agroinfiltration. 0.5 mM SA was applied to plants after agroinfiltration and total protein extract was subjected to immunoblotting using anti-PR1 or anti-GFP antibody.

Given that the chromatin binding of EDS1 was strongly enhanced by SA (Fig. 4a), we next tested whether SA contributes to the nuclear translocation of EDS1 using *35S*:*EDS1-eGFP*/*eds1-2* transgenic plants, which constitutively express EDS1 fused with enhanced GFP (eGFP) in the *eds1-2* background. After SA treatment, we found that neither the accumulation of constitutively expressed EDS1-eGFP protein (Supplementary Fig. 4c) nor its nuclear import (Supplementary Fig. 4d) was apparently induced by SA, in agreement with the finding from another parallel experiment using *35S*:*GFP-EDS1*/*eds1-2* plants (Supplementary Fig. 4e). Thus, SA does not facilitate nuclear translocation of EDS1. It is worthwhile to mention that endogenous EDS1 protein in nuclei is obviously induced by SA (Supplementary Fig. 4f), which is attributed to the fact that the total EDS1 protein expression is enhanced by SA (Figs. 1f and 3a). These data indicate that the accumulation of nuclear EDS1 is required for SA-triggered chromatin binding by EDS1.

### EDS1 functions as a transcriptional coactivator with acidic activation domains

In view of the potential autoactivation of EDS1 observed in Y2H system (Supplementary Fig. 4g), we speculated that EDS1 has transcriptional activator activity. To confirm this, we detected transcriptional activation activity using a yeast monohybrid assay, in which GAL4 DNA binding domain (GAL4 DBD) fusion proteins were transformed into a yeast strain carrying GAL4 promoter-dependent reporter genes^56,57^. Based on *HIS3, MEL1*, and *lacZ* reporter assays (Fig. 4b), we found that full-length EDS1 and TGA3 have transcriptional activation activities in contrast to the GAL4 DBD empty vector, PAD4, and TGA2 (Fig. 4b, top). A more in-depth N-terminal and C-terminal deletion analysis identified two transcriptional activation domains (TADs) located at the α-helical region (331-350) and the C-terminal region (542-593) that are either necessary or sufficient for the transactivation activity of EDS1, respectively (Fig. 4b). Acidic activation domains (AAD), also known as “acidic blobs”, play essential roles in the functions of important transcriptional activators such as p53, GCN4, GAL4, and VP16^58-61^. The acidic amino acids and surrounding hydrophobic residues within AAD have been shown to be critical structural elements for AAD and they are presumably involved in both ionic and hydrophobic interactions with AAD’s target molecules^61^. Importantly, we found that acidic and hydrophobic amino acids are enriched in the TADs of EDS1 (Fig. 4b, bottom), indicating that EDS1 is a transcriptional activator with AADs. Taken together, EDS1 harbors two AADs and has transcriptional activator function.

To further analyze the activator function of EDS1 *in planta*, we investigated the transcriptional regulation of defense genes by EDS1 using *35S*:*EDS1-eGFP*/*eds1-2* transgenic plants. In the transactivation experiments, EDS1-eGFP alone had no effect on gene transcription without SA treatment, but it significantly induced the expression of *PR1* (Fig. 4c) and other defense genes (i.e., *PR2, PR5, PAD4* and *PBS3*) in the presence of SA (Supplementary Fig. 4h). Additionally, EDS1 likely binds to defense gene promoters through intermediate transcriptional regulators owing to lack of a DNA-binding domain^25^. Taken together with the above ChIP and yeast results, these data demonstrate that EDS1 can bind chromatins and acts as a transcriptional coactivator to activate defense genes upon SA induction.

### EDS1 strongly interacts with Mediator

Mediator complex has emerged as a key transcriptional regulator linking different transcription activators and RNA polymerase II (RNAPII) preinitiation complex^62^. CDK8 is a key component in the kinase module (CDK8 module) of the Mediator complex. Increasing studies have demonstrated that CDK8 can play positive roles in gene activation in mammalian and plant cells^63,64^. Of note, we have shown the significant association of plant CDK8 with NPR1 in plants and several components of the CDK8 module positively regulating SA signaling in SAR^65^. To further investigate the mechanism for EDS1-mediated transactivation, we examined the possible association of CDK8 with EDS1 by Co-IP assays. We observed that EDS1 associates with CDK8 in plants (Fig. 4d). To test whether EDS1 physically interacts with CDK8, a reciprocal Y2H assay was used and the direct EDS1-CDK8 interaction was confirmed (Fig. 4e). Moreover, the EDS1 (Δ331-350) deletion mutant is unable to interact with CDK8 (Fig. 4e), suggesting this TAD of EDS1 confers to the interaction with Mediator. These results further support that EDS1 can function as a transcriptional coactivator, which is mediated by interacting with Mediator complex in regulating RNAPII for pathway-specific transcription.

Furthermore, we explored the action of the TAD (331-350) and the NPR1-interacting domain (310-330) of EDS1 (Supplementary Fig. 1b) in defense responses through examining the PR1 expression potentially affected by these EDS1 mutations. Notably, these regions are not highly conserved in PAD4 or SAG101 (Supplementary Fig. 4g). Compared with full-length *EDS1-eGFP*, the accumulation of PR1 was compromised by constructs expressing *EDS1* (Δ310-330)*-eGFP* or *EDS1* (Δ331-350)*-eGFP* driven by a constitutive 35S promoter in Col *eds1-2* plants (Fig. 4f). Therefore, these distinct domains of EDS1 are important for reprogramming gene expression in plant defense.

### EDS1 is directly recruited by NPR1 onto the *PR1* Promoter via a physical NPR1-EDS1 interaction

As shown above, EDS1 and NPR1 occupy the same chromatin loci and synergistically activate plant defense genes (Figs. 2c, 3c and 4a), supporting that EDS1 is a functional NPR1 cofactor in SA-mediated gene regulation. To further explore the effects of SA and NPR1-EDS1 complex on the enrichment of NPR1 and EDS1 at the *PR1* promoter, we performed a series of ChIP and cell fractionation experiments using different transgenic plants constitutively expressing EDS1-eGFP or NPR1-GFP in diverse genetic backgrounds.

First, we determined whether the interaction of NPR1 and EDS1 affected the recruitment of NPR1 and/or EDS1 to the *PR1* promoter. In the ChIP experiments using *35S*:*EDS1-eGFP*/*eds1-2*;*npr1-2* and *35S*:*EDS1-eGFP*/*eds1-2* transgenic lines, *npr1-2* greatly suppressed the occupancy of EDS1-eGFP at specific sites on the *PR1* promoter (Fig. 5a), suggesting that NPR1 is essential for the association of EDS1 with the *PR1* promoter after SA treatment. In contrast, EDS1 appears to only slightly affect NPR1-GFP residence on the *PR1* promoter based on assays using *35S*:*NPR1-GFP*/*npr1-2* and *35S*:*NPR1-GFP*/*npr1-2*;*eds1-2* lines (Supplementary Fig. 5a). These findings indicate that NPR1 is indispensable for EDS1 recruitment at the *PR1* promoter, but not vice versa.

**Fig. 5.**
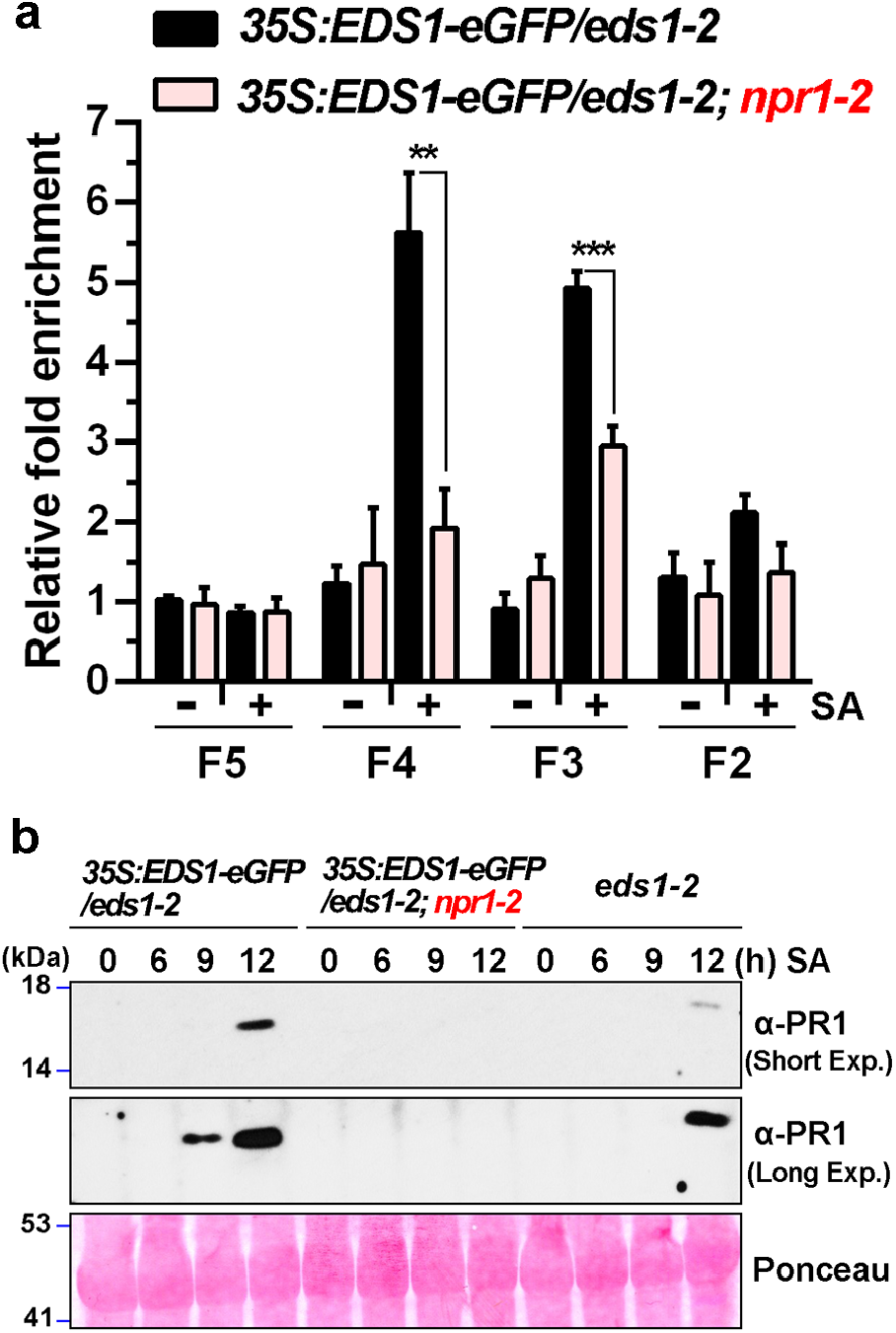
NPR1 directly recruits EDS1 onto promoter and potentiates defense responses. **a**, NPR1 directly recruits EDS1 onto *PR1* promoter. ChIP-qPCR analysis of EDS1-eGFP enrichment at *PR1* genomic loci using indicated soil-grown transgenic plants treated with foliar sprays plus soil-drenches of 0.5 mM SA (+) or water (-) for 9 h. Schematic representation of DNA fragments for amplicons are shown in Fig. 4a. **b**, EDS1 potentiates SA and NPR1-mediated PR1 protein accumulation. Leaves from soil-grown plants were infiltrated with 0.5 mM SA and collected at indicated time points. Short-and long-exposure (Exp.) images of same blot are shown. Total protein was extracted for immunoblotting using an anti-PR1 antibody. Bars indicate SD; n = 3 biologically independent samples. Significances of differences are denoted (*t*-test, **, *P* < 0.01; ***, *P* < 0.001).

Next, we sought to examine the effects of SA and NPR1 on recruiting EDS1 to the *PR1* promoter. Similar to the results obtained with p*EDS1*:*EDS1-FLAG*/*eds1-2* lines (Fig. 4a), the EDS1-eGFP enrichment at the *PR1* promoter in *35S*:*EDS1-eGFP*/*eds1-2* line was also dependent on SA (Fig. 5a). Since SA did not increase the levels of EDS1-eGFP (Supplementary Fig. 4c) or its nuclear translocation (Supplementary Fig. 4d,e), we believe that regulation of nuclear EDS1 by SA is critical for its association with the *PR1* promoter. Notably, our cell fractionation assays demonstrated that neither the EDS1-eGFP expression (Supplementary Fig. 5b) nor its nuclear translocation was promoted by NPR1 after SA treatment (Supplementary Fig. 5c), significantly ruling out the possibility that NPR1 facilitates the nuclear movement of EDS1 upon SA induction. Note that the association of NPR1 with chromatin on the *PR1* promoter was dependent on SA (Fig. 4a). Taken together, these results indicate that SA-induced association of NPR1 with chromatin is crucial for the SA-triggered EDS1 recruitment onto *PR1* promoter, which is independent of the nucleocytoplasmic trafficking of EDS1.

We next focused on investigating the mechanisms utilized by NPR1 in recruiting EDS1 onto the *PR1* promoter. The TGA2 subclade TFs, the major regulators of NPR1-mediated SAR and expression of *PR* genes^51^, are implicated in the recruitment of NPR1 onto the *PR1* promoter^20,66^. NPR1 and EDS1 were enriched at the chromatin site F4 region containing *activation sequence-1* (*as-1*)-like *cis*-elements on the *PR1* promoter (Fig. 4a), an important region for basal and SA-induced *PR1* expression^67^. The SA-responsive *as-1* region is proposed to be occupied by constitutive *trans*-acting factors such as the TGA2/5 and additional factors in the uninduced state^66-68^. Since the constitutively expressed EDS1-eGFP in the nucleus (Supplementary Fig. 4d-f) did not reside at the *PR1* promoter in uninduced states (Fig. 5a), this deduces that EDS1 might not be recruited by the aforementioned *trans*-acting factors. Most importantly, *npr1-2* mutation completely abolished an association of EDS1-eGFP with the F4 (*as-1*) region upon SA induction (Fig. 5a) and EDS1 was not shown to physically interact with TGA2/5 by Y2H assays (Supplementary Fig. 5d), further emphasizing that the direct recruitment of EDS1 onto the *PR1* promoter is predominantly dependent on the physical NPR1-EDS1 interaction.

Based on the above results, we conclude that NPR1 directly recruits EDS1 to the *PR1* promoter, which is crucial for SA-induced EDS1 chromatin binding and *PR1* activation. Consistently, the enhanced activation of *PR1* and other defense genes in the *35S*:*EDS1-eGFP*/*eds1-2* plants upon SA induction is significantly compromised by the *npr1-2* mutation (Fig. 4c and Supplementary Fig. 4h), suggesting that EDS1 cooperates with NPR1 for potentiation of *PR1* expression. This proposition is in line with the immunoblot results showing that NPR1 is required for the accumulation of PR1 protein induced by EDS1-eGFP (Fig. 5b). Overall, EDS1 is directly recruited by NPR1 and participates in transcriptional reprogramming with Mediator complex, and therefore reinforces SA-mediated defense responses.

### NPR1 transcriptionally upregulates *EDS1*

Because EDS1 is apparently induced by SA (Figs. 1f and 3a), we speculate that NPR1 regulates *EDS1* expression. To test this hypothesis, we examined the dynamic expression of EDS1 protein in *npr1-2* and *npr1-3* mutants at different time points. As anticipated, SA-induced EDS1 protein level was obviously reduced in *npr1-2* (Fig. 6a) or *npr1-3* mutants (Supplementary Fig. 6a). SA-upregulated *EDS1* transcript level was also significantly diminished in *npr1-2* (Fig. 6b) or *npr1-3* mutants (Supplementary Fig. 6b), compared with the wild-type control. Therefore, NPR1 preferentially upregulates *EDS1* transcription. Additional ChIP assays demonstrated the SA-dependent association of NPR1-GFP with the *EDS1* promoter at TGA motifs (Fig. 6c, top). Thus, NPR1 directly activates *EDS1* transcription upon SA induction. Since TGA2-NPR1 complex is crucial for *PR1* activation^22,51^, we further test whether the TGA2 also directly targets the *EDS1* promoter. ChIP experiments showed that TGA2-GFP strongly associated with two TGA motifs within the *EDS1* promoter (Fig. 6c, bottom). These data suggest that TGA2-NPR1 complex directly activates *EDS1*.

**Fig. 6.**
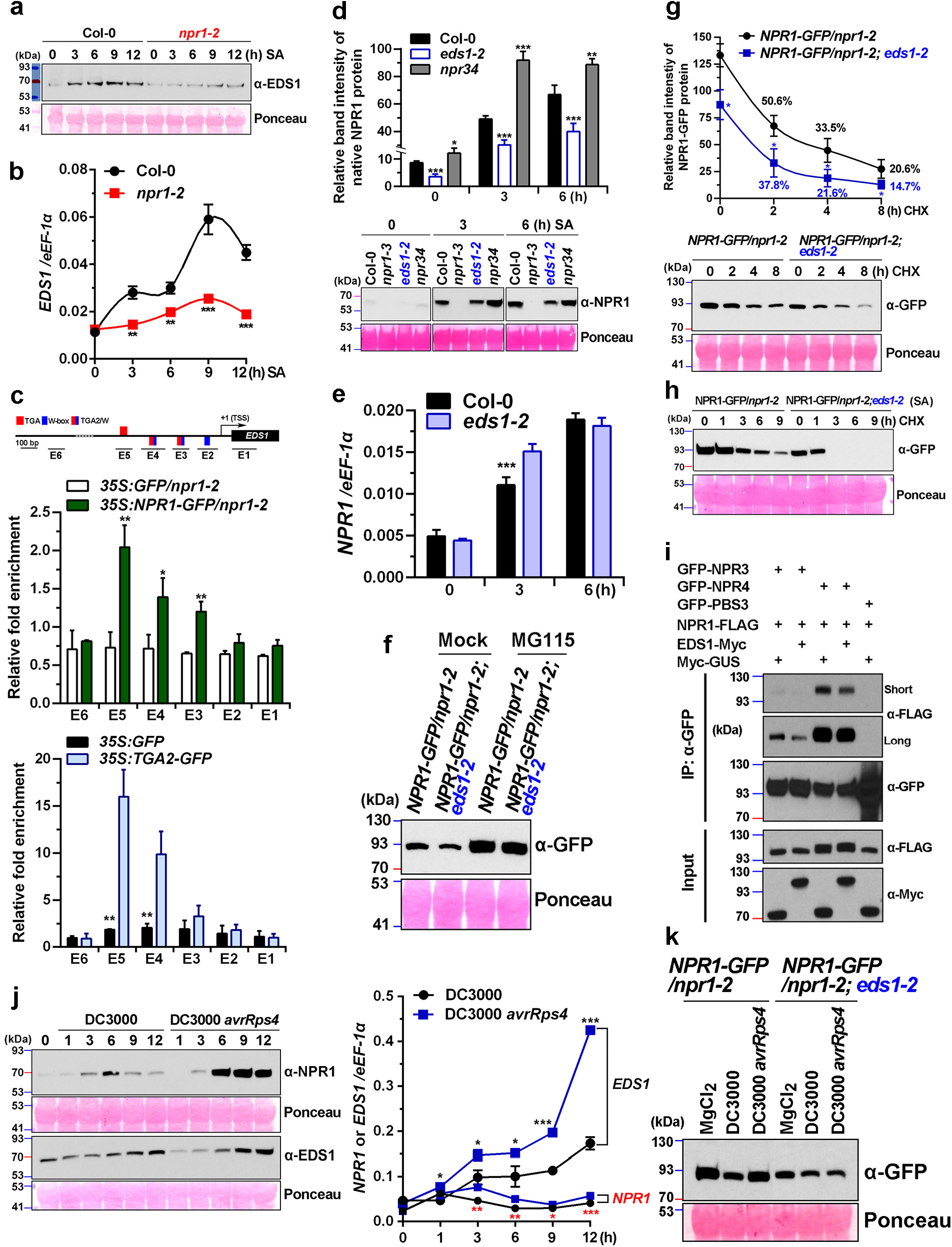
A positive feedback loop of NPR1 and EDS1 in immune responses. **a-c**, NPR1 transcriptionally regulates *EDS1*. EDS1 protein levels (**a**) and *EDS1* mRNA levels (**b**) from seedlings treated with 0.5 mM SA solution for indicated times. ChIP analysis was performed with GFP-Trap magnetic agarose using *NPR1-GFP* transgenic plants treated with 0.5 mM SA for 9 h (**c**). Upper panel, schematic representation of the amplicons in *EDS1* genomic loci. Different *cis*-elements are denoted as described in Fig. 4a. **d-h**, EDS1 stabilizes NPR1. NPR1 protein level (**d**) and *NPR1* mRNA level (**e**) in Col-0 and *eds1-2* seedlings treated with 0.5 mM SA solution for indicated times. NPR1-GFP protein level (**f**,**g)** in *35S*:*NPR1-GFP*/*npr1-2* and *35S*:*NPR1-GFP*/*npr1-2*; *eds1-2* transgenic seedlings treated with 50 µM MG115 or 0.2 mM cycloheximide (CHX) for indicated times. Bars indicate ± SE from three biologically independent experiments (**g**). Percentages indicate the ratio of protein level in CHX treated plants to that of nontreated plants in each genotype. 0.2 mM CHX was infiltrated into rosette leaves of soil-grown plants after treatment with foliar sprays of 0.5 mM SA for 12 h (**h**). **i**, *N. benthamiana* was co-transformed with indicated constructs such as *35S*:*GFP-NPR3*/*4*/*PBS3*, p*NPR1*:*NPR1-3FLAG*, p*EDS1*:*EDS1-9Myc*, and *35S*:*Myc-GUS*. Co-IP assay was performed using GFP-Trap magnetic beads and total protein was analyzed by reducing SDS-PAGE. **j**,**k**, ETI-activated EDS1 protects NPR1 from degradation. NPR1 and EDS1 protein levels (**j**, left) and their corresponding mRNA levels (**j**, right) in Col-0 plants infiltrated with *Pst* DC3000 or *Pst* DC3000 *avrRps4* (OD_600_ = 0.01) for indicated times. NPR1-GFP protein level (**k**) in 4-week-old soil-grown *35S*:*NPR1-GFP*/*npr1-2* and *35S*:*NPR1-GFP*/*npr1-2*;*eds1-2* transgenic plants infiltrated with MgCl_2_or pathogens (OD_600_= 0.01). Bars indicate SD. n = 3 biologically independent samples (**b**,**c**,**j**); n = 4 biologically independent samples (**d**,**e**). Significances of differences from the control are shown for each time point or each amplicon (*t*-test, *, *P* < 0.05; **, *P* < 0.01; ***, *P* < 0.001). Total RNA was extracted from 2-week-old seedlings and subjected to qPCR analysis. Total protein was analyzed by reducing SDS-PAGE and immunoblotting using indicated antibodies.

### EDS1 Protein Stabilizes NPR1 Protein

In *Arabidopsis*, NPR1 protein is ubiquitously turned over by proteasome-mediated protein degradation and moderately regulated by gene regulation^33,55,65^. We asked whether EDS1 regulates NPR1 at the transcriptional or translational level. In the dynamic expression assays, the basal and SA-induced NPR1 protein levels were significantly reduced in *eds1-2*, compared with wild-type control (Fig. 6d and Supplementary Fig. 6c,e). However, *NPR1* transcript level in *eds1-2* was comparable to that in control plants (Fig. 6e and Supplementary Fig. 6d), albeit unexpectedly increased at 3 h after SA treatment. These results strongly indicate that EDS1 post-transcriptionally regulates NPR1. To further investigate whether EDS1 stabilizes the NPR1 protein, we compared protein levels of the constitutively expressed NPR1-GFP in *35S*:*NPR1-GFP*/*npr1-2* and *35S*:*NPR1-GFP*/*npr1-2*;*eds1-2* transgenic lines. In *35S*:*NPR1-GFP*/*npr1-2*;*eds1-2* plants, there were reduced levels of NPR1-GFP protein, which could be restored in the presence of the 26S proteasome inhibitor MG115 (Fig. 6f). We then analyzed EDS1 protein stability using cycloheximide, a potent protein synthesis inhibitor. In a full cycloheximide-chase assay, *eds1-2* mutation significantly accelerated the decay of uninduced NPR1-GFP protein (Fig. 6g) and strongly improve the decay of SA-induced NPR1-GFP protein (Fig. 6h). In contrast, *npr1-2* mutation did not affect cycloheximide-resistant GFP-EDS1 or EDS1-eGFP (Supplementary Fig. 6f,g), consistent with above finding that NPR1 preferentially regulates *EDS1* transcription. These results provide compelling evidence that EDS1 stabilizes NPR1 in order to maintain an optimal NPR1 threshold for plant defense responses.

The SA receptors NPR3 and NPR4 were proposed to be the adaptors of a Cullin3-based E3 ligase and promote the degradation of NPR1, EDS1, and JAZ proteins^17,69,70^. We further examined the biochemical basis underlying the NPR1 homeostasis regulated by EDS1. In Co-IP assays, GFP-NPR3 and GFP-NPR4 were efficiently immunoprecipitated with NPR1-FLAG (Fig. 6i and Supplementary Fig. 6h). However, expression of EDS1-Myc protein diminished the amount of NPR1-FLAG bound to GFP-NPR3/NPR4 (Fig. 6i). These results indicate that EDS1 competes with NPR3/NPR4 for NPR1 interaction, thereby stabilizing NPR1 in plant cells.

### NPR1 is stabilized by EDS1 during ETI to confer a robust defense

To decipher the mechanism underlying the regulation of NPR1 stability in plant-pathogen interactions, we first investigated the accumulation of NPR1 protein in response to virulent and avirulent pathogen challenges. Time-course expression analyses showed that infection with avirulent *Pst* DC3000 *avrRps4* induced NPR1 protein more strongly than inoculation by virulent *Pst* DC3000 (Fig. 6j, left), which obviously differs from the gene transcription patterns (Fig. 6j, right). These results demonstrate that NPR1 protein rather than its transcript hyperaccumulates during RRS1S/RPS4-activated ETI, suggesting that ETI preserves NPR1.

Although ETI slightly enhanced the induction of *NPR1* transcription (Fig. 6j, right) to compensate for the degradation of NPR1 promoted by *Pst* DC3000 in *35S*:*NPR1-GFP*/*npr1-2* transgenic plants (Fig. 6k), it is reasonable to speculate that ETI prevents NPR1 degradation. As anticipated, we found that the destruction of NPR1-GFP protein caused by *Pst* DC3000, as reported previously^34^, was apparently restored by *Pst* DC3000 *avrRps4* (Fig. 6k), further supporting that ETI prevents NPR1 degradation. Moreover, the *eds1-2* mutation impeded the recovery of NPR1-GFP protein by *Pst* DC3000 *avrRps4* in *35S*:*NPR1-GFP*/*npr1-2*;*eds1-2* lines (Fig. 6k), indicating the prevention of NPR1 degradation by ETI occurred in an EDS1-dependent manner. Consistent with the above results, these findings confirm that EDS1 protects NPR1 from degradation in plant-pathogen interactions.

## Discussion

Genetic studies have identified several important positive regulators of plant immunity, including NPR1 (ref.^38^), EDS1 (ref.^25^), PAD4 (ref.^40^), NDR1 (ref.^71^), PBS3 (ref.^72^), EDS5 (ref.^73^), and EPS1 (ref.^74^). Among them, the transcriptional regulator NPR1 has been known as the master regulator of SA signaling and SAR^13,41,75^. However, the mechanisms of NPR1-mediated transcriptional reprogramming are still poorly understood. EDS1 is required for plant basal defense, TIR-NLR-mediated ETI and SAR^25,28^ and regulates the expression of a large number of defense-related genes^31^. Nonetheless, how EDS1 activates downstream plant defense genes remains obscure.

In the present study, we have shown the functionally physical and genetic interactions between two key immune regulators for the synergistical control of plant immune responses. The proposed model is illustrated in the Supplementary Fig. 6i. We provide the first evidence suggesting that EDS1 is capable of acting as a transcriptional coactivator, which cooperates with NPR1 and Mediator in the transcription machinery for enhancing activation of defense genes upon immune induction. Interaction between two coactivators promotes direct recruitment of EDS1 onto promoters and influences the homeostasis of protein by stabilizing NPR1. We have elucidated an elaborate positive-feedback regulation of NPR1 and EDS1 by distinct mechanisms for amplifying defense responses. EDS1 was classified as a lipase-like protein^25,29^, but subsequent biochemical and structural studies showed that EDS1 has no lipase activity^29,76^. Our study demonstrates that EDS1 serves as a transcriptional coactivator based on the following criteria. First, EDS1 binds chromatin regions in a stimulus-specific manner (Fig. 4a and Supplementary Fig. 4a,b) and directly interacts with the transcriptional coactivator NPR1 (Fig. 1a-f). Second, EDS1 possesses transactivation activity and contains two intrinsic TADs (Fig. 4b). We further found that acidic and hydrophobic amino acids are overrepresented within EDS1’s two discrete TADs (Fig. 4b, bottom), indicating that EDS1 is a transcriptional activator with AADs. In addition, EDS1 activates many defense genes in response to SA (Fig. 4c and Supplementary Figs. 3a-c and 4h). Furthermore, one TAD of EDS1 directly interacts with a subunit of the Mediator (i.e., CDK8) (Fig. 4d,e), further indicating that EDS1 is a transcriptional activator that recruits Mediator complex in the transcription machinery. Since EDS1 itself likely does not bind chromosomal DNA directly^25^, these findings strongly support that EDS1 is a bona fide transcriptional coactivator. It is worthwhile to mention that all previously reported AADs were identified in transcription activators with a DNA binding domain^57-60^. For the first time, we have shown that EDS1, a transcriptional coactivator without a DNA binding domain, possesses two discrete AADs. Therefore, our study may shed light on the functions of transcriptional coactivators in general. Mechanistically, EDS1 is directly recruited to the specific SA-responsive *cis*-elements on *PR1* promoter by NPR1 and works together with NPR1, thus enhancing *PR1* transcription in SA-mediated defense (Fig. 5a,b and Supplementary Fig. 6i). Interestingly, apart from associating with TGA motif-containing chromatin regions, EDS1 has also been shown to reside at the promoter regions containing W-box *cis*-elements in a similar manner to NPR1 (Fig. 4a and Supplementary Fig. 4a,b). Consistently, several WRKY factors have been shown to interact with NPR1^65,77^. Given that multiple WRKY factors exhibit intricate redundancy, cooperation, and antagonism on gene regulation and disease resistance to different pathogens^53,54^, the interaction of the NPR1-EDS1 complex with diverse TFs might fine-tune the dynamic gene expression regulating plant growth and immune responses.

The interaction between NPR1 and the TGA2 subclass of TFs has been shown to play an important role in activating plant defense gene expression^22,66^. It is suggested that NPR1 is recruited by TGA2 onto the *PR1* promoter upon SA induction^20,66^, but whether NPR1 directly recruits transcriptional (co)factors to promote defense gene expression remains unknown. Our data indicate that NPR1 directly recruits a novel transcriptional coactivator EDS1 onto the *PR1* promoter via a physical interaction to stimulate *PR1* expression (Figs. 5a,b). Therefore, these findings suggest a novel prominent regulatory role of NPR1’s transactivation, which is required for mediating the assembly of multiple regulatory activators for specific transactivation.

Intriguingly, SA treatment activates the transactivation function of NPR1 presumably by releasing the autoinhibition of its cryptic transactivation activity^22^. This relief of repression model suggests that NPR1 can act as a coactivator in an SA-dependent manner. Our study implies that the dynamic interaction of NPR1 with certain SA-induced regulators (e.g., EDS1) (Supplementary Fig. 1b-e) may contribute to the relief of the repression of NPR1’s transactivation activity by inducing conformation changes. Further structural and biochemical analyses of NPR1 and its partners are needed to test this possibility.

In mammalian systems, several studies have shown that diverse endogenous transcriptional activators form transcriptional condensates with the Mediator complex to robustly drive gene expression^78,79^. Our study shows that two interacting transcriptional coactivators form nuclear foci in plant cell nuclei and interact with a component of the Mediator, CDK8 (Figs. 1e and 4d,e). Thus, these plant coactivators may form phase-separated nuclear condensates for active transcription. Most recently, NPR1 has been reported to facilitate the formation of cytoplasmic condensates for degradation of substrates to inhibit cell death^80^. Nonetheless, nuclear NPR1 and EDS1 may incorporate diverse transcriptional (co)factors into transcriptional activator concentrates for robust transcriptional reprograming to relocate energy for defense instead of growth upon pathogen infection.

EDS1-mediated signaling can boost SA accumulation (upstream of SA) in innate immunity^46^. In this study, we further demonstrate that EDS1 also acts as an essential positive regulator of SA signaling (downstream of SA) because it significantly facilitates expression of *PR* and other defense genes in response to SA (Fig. 3a and Supplementary Fig. 3a-c). Furthermore, EDS1 and NPR1 synergistically accelerate transcriptional reprograming and promote pathogen resistance (Fig. 3b,c and Supplementary Fig. 3d). Consistently, EDS1-mediated SA signaling rather than SA accumulation is able to contribute to RRS1S/RPS4-mediated ETI, because overexpression of *EDS1* results in enhanced responsiveness to exogenous SA for protection against pathogen^81^. Consequently, EDS1 functions both upstream and downstream of SA for SA-mediated defense, which is similar as *Arabidopsis* ELP2, an accelerator of immune responses^82^. Thus, some immune regulators potentiate plant defense through promoting both signaling transduction and biosynthesis of SA.

This study shows that EDS1 rather than its partners (i.e., PAD4 and SAG101) possesses intrinsic transactivation activity (Fig. 4b and Supplementary Fig. 4g). As PAD4 and SAG101 are required for accumulation of EDS1 (ref.^76^) and as no direct interaction of NPR1 with PAD4 or SAG101 is detected (Fig. 1a and Supplementary Fig. 1a), it is suggested that EDS1-PAD4 and/or EDS1-SAG101 complex may preferentially contribute to NPR1-mediated gene activation by stabilizing EDS1 protein in plant-pathogen interactions. On the other hand, nucleocytoplasmic coordination of EDS1 and its interacting factors are involved in cell compartment-specific and full immune responses^27,45^. NPR1, EDS1, and PAD4 are localized in the cytosol and nucleus^18,19,76^, while SAG101 is exclusively detected in the nucleus^76^. It is suggested that neither SAG101 nor PAD4 affects nucleocytoplasmic localization of EDS1^76^. However, nucleocytoplasmic EDS1-PAD4 is required for signal transduction in basal immunity and SAR^28,76^; nuclear EDS1-SAG101 may be important for nuclear EDS1 retention^76^. In this work, nucleocytoplasmic NPR1 does not affect the intracellular trafficking of EDS1 from cytoplasm to nucleus (Supplementary Fig. 5c). Instead, nuclear EDS1-NPR1 association is markedly enhanced in the specific nuclear compartment, and in turn contributes to chromatin binding of EDS1 under induced states (Figs. 1e,f and 5a). Thus, the intricate dynamic association of EDS1 with its partners are essential for temporal and spatial coordination of diverse immune responses.

Multiple lines of evidence indicate that EDS1 control plant immunity during diverse pathways^28,31^. Our study primarily shows the functions of nuclear NPR1-EDS1 association with chromatin in SA signaling, whereas EDS1-PAD4 has been shown to work redundantly with SA at early defense signaling^83-85^. A reciprocal antagonism between SA and jasmonic acid (JA)-regulated transduction pathways play key roles in resistance to diverse pathogens^86^. NPR1 has been reported to inhibit JA signaling by suppressing JA-responsive gene expression^87^, while EDS1-PAD4 antagonizes MYC2-mediated JA signaling for RRS1S/RPS4-mediated immunity^84^. It seems that both NPR1 and EDS1 play a role in the crosstalk between SA and JA-mediated pathways. More studies are needed to further dissect the mechanisms of SA and JA crosstalk regulated by the interplay of NPR1 and EDS1.

In summary, this work sheds light on the function of a novel transcriptional coactivator complex at the epicenter of plant immunity. Our study has revealed uncharacterized roles of NPR1 and EDS1 in signal transduction and activation of immune responses. Identification of EDS1 as a novel transcriptional coactivator not only opens a new avenue for studying the signaling pathways in plant immune responses, but also sheds light on the molecular basis for general gene regulation. Meanwhile, direct recruitment of coactivator by NPR1 upon immune induction provides new insight into the mechanism of NPR1’s transactivation.

## Methods

### Plant materials and growth conditions

*Arabidopsis thaliana* (L.) Heynh. seeds were sown on autoclaved soil and vernalized at 4°C for 3 days. Plants were germinated and grown in a growth chamber at 22°C day/20°C night with ∼70% relative humidity and 12-h light/12-h dark photoperiod for middle-day conditions. To grow *Arabidopsis* seedlings *in vitro*, seeds were first sterilized by chlorine gas for 3 h in a desiccator and sown on sterilized half-strength MS media (pH 5.7) supplemented with 1% sucrose and 0.25% phytagel with appropriate antibiotics. Plated seeds were stratified at 4°C for 3 days and then germinated in a growth chamber at 22°C day/20°C night under 16-h light/8-h dark photoperiod for long-day conditions. *N. benthamiana* was grown in a growth chamber at 25°C under middle-day conditions.

The *npr1-2* (ref.^13^), *npr1-3* (ref.^40^), *eds1-2* (ref.^31^), *pad4-1* (ref.^40^), *rps4-2* (ref.^88^) and *npr3-2 npr4-2* (ref.^17^) mutants are in the Columbia (Col-0) ecotype. The *eds1-1* (ref.^25^) is in the Wassilewskija (Ws-0) ecotype.

### Constructs, transgenic plants and genetic analysis

For generating expression constructs, the Gateway Cloning Technology (Invitrogen) and In-Fusion Advantage PCR Cloning Kit (Clontech) were used. Most DNA fragments were amplified and cloned into entry vectors such as pDONR207 and pENTR/D-TOPO (Invitrogen), and then transferred to the destination vectors. The binary vectors were transformed into *Agrobacterium* by electroporation and then transformed into *N. benthamiana* or *Arabidopsis* lines. The stable T_2_ transgenic lines with single inserts were analyzed and carried to produce T_3_ homozygous progenies. At least two independent homozygous lines expressing target protein significantly were selected for further studies in all experiments. The primers (Supplementary Table 1) and recombinant DNA constructs (Supplementary Table 2) for all experiments are listed and described previously^34^.

To create p*EDS1*:*EDS1-FLAG* expression clones, the genomic coding region and 2-kb upstream sequences of *EDS1* DNA was cloned into entry clone and then transferred to the pEarleyGate302-3xFLAG destination vector kindly provided by Xuehua Zhong (University of Wisconsin-Madison); the combined binary vector was introduced into *Arabidopsis eds1-2* mutant background to obtain the p*EDS1*:*EDS1-FLAG*/*eds1-2* transgenic lines. For *35S*:*EDS1-eGFP*/*eds1-2* and *35S*:*GFP-EDS1*/*eds1-2* transgenic lines, the full-length EDS1 cDNA was cloned into entry vector and transferred into pK7FWG2 and pMDC43 destination vector; these binary plasmids were introduced into *eds1-2* plants, respectively. To generate *35S*:*GFP-NPR1* overexpression transgenic lines and *35S*:*NPR1-GFP*/*npr1-2* plants, full-length NPR1 cDNA was cloned into pMDC43 destination vector and pCB302 binary vector and then the resulting vectors were introduced into Col-0 wild-type and *npr1-2* mutant backgrounds, respectively.

All crosses among different genotypes were performed by pollinating the emasculated flowers of maternal recipient with pollen from male donor. The *npr1-2 eds1-2* and *eds1-2 npr1-2* double mutants were generated by crossing female *npr1-2* with *eds1-2* and by crossing female *eds1-2* with *npr1-2*, respectively. To generate *35S*:*EDS1-eGFP*/*eds1-2*; *npr1-2* lines, *npr1-2* was crossed with *35S*:*EDS1-eGFP*/*eds1-2* plants. For *35S*:*NPR1-GFP*/*npr1-2*; *eds1-2* lines, *35S*:*NPR1-GFP*/*npr1-2* as a recipient was crossed with *eds1-2*. The double mutations in the segregating F_2_ populations was identified by a *npr1-2* CAPS maker and by PCR using primers flanking the *eds1-2* deletion region; the homozygosity for the EDS1-eGFP or NPR1-GFP transgene was confirmed in next generation by genotyping using specific primers for eGFP or GFP. All the successive plants and controls at the same generation were selected in further study. To generate p*NPR1*:*Myc-NPR1* plants containing p*EDS1*:*EDS1-FLAG*, the p*EDS1*:*EDS1-FLAG*/*eds1-2* transgenic plants was crossed with p*NPR1*:*Myc-NPR1/npr1-3* plants^89^provided from Zhonglin Mou (University of Florida). The F_2_ plants were selected on antibiotics and genotyped using *npr1-3* CAPS marker^82^ and specific primers for *eds1-2* and transgene.

### Y2H and yeast monohybrid assays

Y2H assays were performed as described previously^34^. The pDEST-GBKT7 based bait vectors were transformed into the yeast strain Y187 and the yeast strain AH109 was transformed with pDEST-GADT7 based vectors. The fresh diploids by yeast mating were used to detect protein-protein interactions on selective media.

For yeast monohybrid assays, pDEST-GBKT7 based GAL4 DNA-BD fusion vectors were transformed into the yeast strain AH109 including several reporters (*HIS3, MEL1* and *lacZ*) under distinct GAL4 upstream activating sequences as described in Matchmaker GAL4 Two-Hybrid System 3 & Libraries User Manual (Clontech). The transformants were grown on synthetic dropout (SD) agar medium lacking Trp and His (-WH) and detected on SD/-WH/ X-α-Gal (Biosynth). The liquid cultures of yeast cells were used to detect the *lacZ* expression in quantitative β-galactosidase assays with o-Nitrophenyl-β-D-Galactopyranoside (ONPG, Amresco) performed according to the Yeast Protocols Handbook (Clontech).

### Pull-down assay

The recombinant protein expression and in vitro pull-down assay were carried out as previously described^34^ with minor modification. For GST-fusion protein expression, the coding sequences of GUS, EDS1 and PAD4 were cloned into entry clone and transferred into pDEST15, respectively. These GST-fusion constructs and GST (empty pGEX-4T-1 vector) were heterologously expressed in the *E. coli* Rosetta (DE3) cell line. The Trx-His_6_-NPR1 protein was expressed in expressed in *E. coli* OverExpress™ C41 (DE3) strain using the plasmid pET-32a. For the pull-down assay, Trx-His_6_-NPR1 protein in 2 ml of extracts was immobilized on 30 µL Ni-NTA agarose at 4°C for 1 h. After washed for several times, the whole cell extract of GST-protein fusion was added to each immobilized sample for 1 h at 4°C. After washing, the bound proteins were eluted by boiling in sample buffer and subjected to immunoblotting analysis. The signals were visualized as described previously.

### *Agrobacterium*-mediated transient expression

*Agrobacterium*-mediated transient expression in *N. benthamiana* leaves were performed as described previously^34^. *A. tumefaciens* strain (GV3101/PMP90) carrying the indicated constructs were used together with the p19 strain for infiltration of 2∼4-week-old *N. benthamiana* leaves using a needleless syringe. For transient expression in *Arabidopsis*, approximately 10 young leaves of *eds1-2* mutant were infiltrated with *A. tumefaciens* strain (AGL) containing each construct according to a described method^90^. After agroinfiltration with the presence of 0.01% Silwet L-77, plants were immediately covered, kept in the dark for 24 h and subsequently incubated under middle-day conditions for another 2∼3 days.

### BiLC and BiFC assays

For BiLC assay, the full-length coding sequence of target gene was fused to the N or C terminus of firefly luciferase using pCAMBIA1300 nLUC or pCAMBIA1300 cLUC vector; SGT1b-nLUC and cLUC-RAR1 constructs were used a positive interaction control^91^. Leaves excised 2 days after transient expression were sprayed with luciferin solution (100 µM luciferin and 0.01% Triton X-100) and kept in the dark for 2 h to quench fluorescence. Luc activity was observed with a low-light cooled CCD imaging apparatus (Andor iXon). In BiFC assay, the relative entry clone was transferred into pMDC43-nVenus and pMDC43-cCFP vectors^26^ provided by Walter Gassmann (University of Missouri). The leaf tissues from the infiltrated area were observed under a confocal microscope (Leica TCS SP8) with the VENUS/GFP filters: 488 nm excitation and 530 nm emission.

### Co-IP and immunoblotting assays

Protein fractionations for immunoblotting and Co-IP assays in *N. benthamiana* and *Arabidopsis* were performed as previously described^34^. The homogenate was sonicated on ice and optionally treated with Benzonase Nuclease (MilliporeSigma) for 30 min on ice. The solution was filtered through Miracloth (Calbiochem). The Myc-Trap^®^_MA (Chromotek) and anti-FLAG^®^ M2 magnetic beads (Sigma-Aldrich) were used to immunoprecipitated the protein complexes. Immunoblotting was performed with anti-Myc Tag (ThermoFisher) and anti-FLAG^®^ M2 antibodies (Sigma-Aldrich).

### Pathogen growth assays

Inoculation of plants with pathogens and pathogenicity tests were performed as described previously^34^. Three full-grown leaves on each 4∼6-week old plant grown under middle-day conditions were inoculated with different *Pseudomonas* strains. The three leaf discs from individual plant were pooled for each sample and six such replicates were used for each genotype in pathogen growth assay.

### Real-time quantitative PCR

Gene expression analysis by qPCR was carried out as previously described ^34^ with minor modification. Total RNA was extracted using RNAzol^®^RT (Sigma-Aldrich) and 2 µg of total RNA was subjected to reverse transcription using qScript cDNA Synthesis Kit (Quanta). Real-time PCR was performed using PerfeCTa SYBR Green FastMix (Quanta). The primers used for qPCR in this study are shown in Supplementary Table 1.

### Cell fractionation

Preparation of nuclear and cytoplasmic fractions was performed according to the user manual supplied with the CelLytic™ PN Plant Nuclei Isolation/Extraction Kit (Sigma) with minor modifications. Approximately 2 g of plant tissues were suspended in nuclei isolation buffer (NIB) and passed through a provided filter mesh. After centrifugation for 15 min, the supernatant was used for further extraction of cytoplasmic proteins and the pellet was used to further extract nuclei and nuclear proteins. The transferred supernatant was centrifugated for 10 min at 12,000 rpm, 4°C and the clean supernatant was collected as cytoplasmic fractions. The initial pelleted nuclei were resuspended in 10 ml NIBA (1X NIB, 1 mM DTT, 1X protease inhibitor cocktail and 0.5% Triton X-100). After centrifugation, isolation of nuclei was carried out as described with Semi-pure Preparation of Nuclei Procedures based on the manufacture protocol. The cellular fractions were analyzed on reducing SDS-PAGE and transferred to Nitrocellulose membranes. PEPC and RuBisCo were detected and used as cytoplasmic markers, and histone H3 was used as nuclear marker.

### ChIP analysis

ChIP was performed according to a previous report^92^ with modifications. Approximately 3 g of 4-week-old soil-grown plants or 3-week-old seedlings were harvested and vacuum infiltrated with 1% formaldehyde for cross-linking. The cross-linking reaction was subsequently stopped by 150 mM glycine. Samples were washed three times with sterile deionized water, dried on paper towel, frozen and stored at -80°C for further use. For chromatin isolation, plant tissues were ground to a fine powder in liquid nitrogen and mixed with 30 ml cold nuclei isolation buffer (0.25 M sucrose, 15 mM PIPES pH 6.8 or 10 mM Tris-HCl pH 7.5, 5 mM MgCl_2_, 60 mM KCl, 15 mM NaCl, 1 mM CaCl_2_, 1% Triton X-100, 1 mM PMSF, 2 µg/ml pepstatin A, 2 µg/ml aprotinin, and 1 mM DTT). Samples were incubated on ice for 5 min with gentle vortex and then filter through two layers of Miracloth (Calbiochem) and centrifuged at 4°C, 3000 *g*, for 20 min. The nuclear pellets were gently resuspended in 1.5 ml of cold nuclei lysis buffer (50 mM Tris-HCl pH 7.5, 150 mM NaCl, 1 mM EDTA, 0.3% sarkosyl, 1% Triton X-100, 50 µM MG-115, 1 mM PMSF, protease inhibitor cocktail, and 1 mM DTT) and incubated on ice with gentle mixing for 5 min. Chromatin was sheared into approximate 500 bp DNA fragments using M220 Focused-ultrasonicator (Covaris) and centrifuged at 13,000 *g*, 4°C for 15 min. The supernatant was collected for further steps. For the immunoprecipitation step, the samples were diluted with ChIP dilution buffer (20 mM Tris-HCl pH 7.5, 150 mM NaCl, 1 mM EDTA, 1% Triton X-100, 50 µM MG-115, 1 mM PMSF, protease inhibitor cocktail, and 1 mM DTT) and precleared for 1 h using control magnetic agarose beads blocked with 100 µg/µl BSA. After removing the beads, 5% of precleared chromatin was retained as input control. Meanwhile, the remaining samples were mixed with conjugated anti-Myc tag antibody (Abcam) with Magna ChIP™ Protein A Magnetic Beads (Sigma-Aldrich) for Myc-NPR1, anti-FLAG^®^ M2 magnetic beads (Sigma-Aldrich) for FLAG-EDS1, or GFP-Trap^®^_MA beads (Chromotek) for EDS1-eGFP. The mixture was incubated at 4°C for 4 h with gentle rotation and then the immunocomplexes were washed twice each with low salt, high salt, LiCl, and TE buffer.

In the reverse cross-linking steps, ChIP sample and input control were mixed with 20% Chelex^®^ 100 Resin (Sigma-Aldrich) solution at room temperature and incubated for 10 min at 95°C shaking every 3 min. Once the sample was cooled down, 20 µg of proteinase K (Invitrogen) was added to a final volume of 200 µl of ChIP reaction in TE, and incubated at 50°C for 1 h followed by boiling for 10 min. After spin down, the supernatant was transferred and retained, the pelleted beads were washed with TE; the washing flow-through was added to the initial supernatant. Then 5 µg of RNase A (Thermo Scientific) was added into each sample and incubated at 37°C for 30 min. Immunoprecipitated DNA was purified using a mixture of phenol:chloroform:isoamyl alcohol (25:24:1) followed by chloroform extraction, ethanol precipitated using Dr. GenTLE Precipitation Carrier (TaKaRa) with incubation at -80°C for 1 h, recovered by centrifugation, washed and resuspended in 100 µl of TE.

Recovered DNA was quantified by qPCR described as above, with the locus-specific primers (Supplementary Table 1) and ChIP-qPCR was performed with at least three technical replicates. Relative DNA level for each amplicon was calculated against the total input using the ΔΔCt_T_ method. Relative fold enrichment was standardized to the *Actin2* open reading frame.

### Quantification and statistical analyses

The results of Western Blots were quantified with software ImageJ (NIH). Statistical analysis was conducted with the software of GraphPad Prism 6.0 using one-way or two-way analysis of variance (ANOVA) with Tukey’s multiple comparisons test or using multiple Student’s *t* tests. Error bars represent standard deviations (SD) or standard errors (SE). Statistically significant differences are marked with asterisks (*t*-test, * *P* < 0.05; **, *P* < 0.01; ***, *P* < 0.001) or different letters (*P* < 0.05). For instance, different letter (A, B, C, etc) are used to label samples with statistical differences, whereas the “ABC” is used to mark samples with no statistical difference to other samples labeled with “A”, “B” or “C”. Detail statistical differences can be found in the figures and figure legends.

### Reporting Summary

Further information on research design is available in the Nature Research Reporting Summary linked to this article.

## Data availability

All supporting data are available in the main text, Supplementary Figs. 1-6 and Supplementary Tables 1-2 in the Supplementary Information. Source data are provided with this paper. Any additional data that support the findings of this study are available from the corresponding authors upon reasonable request. The databases that we used are all publicly available.

## Acknowledgments

We are grateful to Xinnian Dong for *35S*:*NPR1(nls)-GFP* transgenic line, Zhonglin Mou for p*NPR1*:*Myc-NPR1* transgenic line, Walter Gassmann for BiFC constructs and Xuehua Zhong for pEarleyGate302-3xFLAG plasmid. We thank Beth Krizek and David Reisman for critical reading of this manuscript.

## Author contributions

H.C., F.L., and Z.Q.F conceived and designed the experiments. H.C. performed most experiments with assistance with M.L., G.Q., M.Z., L.L., and J.Z.. All authors participated in results discussion and data analysis. H.C., F.L., and Z.Q.F wrote the manuscript with contribution from D.W.

## Additional information

### Supplementary information

is available for this paper.

### Correspondence and requests for materials

should be addressed to F.L. or Z.Q.F.

## References

1. Dodds, P.N. & Rathjen, J.P. Plant immunity: towards an integrated view of plant-pathogen interactions. Nat Rev Genet 11, 539–548 (2010).

2. Dangl, J.L., Horvath, D.M. & Staskawicz, B.J. Pivoting the plant immune system from dissection to deployment. Science 341, 746–751 (2013).

3. Wan, L. et al. TIR domains of plant immune receptors are NAD(+)-cleaving enzymes that promote cell death. Science 365, 799–803 (2019).

4. Burdett, H. et al. The Plant “Resistosome”: Structural Insights into Immune Signaling. Cell Host Microbe 26, 193–201 (2019).

5. Hartmann, M. & Zeier, J. N-hydroxypipecolic acid and salicylic acid: a metabolic duo for systemic acquired resistance. Curr Opin Plant Biol 50, 44–57 (2019).

6. Fu, Z.Q. & Dong, X. Systemic acquired resistance: turning local infection into global defense. Annu Rev Plant Biol 64, 839–863 (2013).

7. Pieterse, C.M., Van der Does, D., Zamioudis, C., Leon-Reyes, A. & Van Wees, S.C. Hormonal modulation of plant immunity. Annu Rev Cell Dev Biol 28, 489–521 (2012).

8. An, C.F. & Mou, Z.L. Salicylic Acid and its Function in Plant Immunity. Journal of Integrative Plant Biology 53, 412–428 (2011).

9. Mishina, T.E. & Zeier, J. Pathogen-associated molecular pattern recognition rather than development of tissue necrosis contributes to bacterial induction of systemic acquired resistance in Arabidopsis. Plant J 50, 500–513 (2007).

10. Wildermuth, M.C., Dewdney, J., Wu, G. & Ausubel, F.M. Isochorismate synthase is required to synthesize salicylic acid for plant defence. Nature 414, 562–565 (2001).

11. Ryals, J.A. et al. Systemic acquired resistance. Plant Cell 8, 1809–1819 (1996).

12. Durner, J., Shah, J. & Klessig, D.F. Salicylic acid and disease resistance in plants. Trends in Plant Science 2, 266–274 (1997).

13. Cao, H., Glazebrook, J., Clarke, J.D., Volko, S. & Dong, X. The Arabidopsis NPR1 gene that controls systemic acquired resistance encodes a novel protein containing ankyrin repeats. Cell 88, 57–63 (1997).

14. Wu, Y. et al. The Arabidopsis NPR1 protein is a receptor for the plant defense hormone salicylic acid. Cell Rep 1, 639–647 (2012).

15. Manohar, M. et al. Identification of multiple salicylic acid-binding proteins using two high throughput screens. Front Plant Sci 5, 777 (2014).

16. Ding, Y. et al. Opposite Roles of Salicylic Acid Receptors NPR1 and NPR3/NPR4 in Transcriptional Regulation of Plant Immunity. Cell 173, 1454–1467 e1415 (2018).

17. Fu, Z.Q. et al. NPR3 and NPR4 are receptors for the immune signal salicylic acid in plants. Nature 486, 228–232 (2012).

18. Mou, Z., Fan, W. & Dong, X. Inducers of plant systemic acquired resistance regulate NPR1 function through redox changes. Cell 113, 935–944 (2003).

19. Kinkema, M., Fan, W. & Dong, X. Nuclear localization of NPR1 is required for activation of PR gene expression. Plant Cell 12, 2339–2350 (2000).

20. Fan, W. & Dong, X. In vivo interaction between NPR1 and transcription factor TGA2 leads to salicylic acid-mediated gene activation in Arabidopsis. Plant Cell 14, 1377–1389 (2002).

21. Li, M. et al. TCP Transcription Factors Interact With NPR1 and Contribute Redundantly to Systemic Acquired Resistance. Front Plant Sci 9, 1153 (2018).

22. Rochon, A., Boyle, P., Wignes, T., Fobert, P.R. & Despres, C. The coactivator function of Arabidopsis NPR1 requires the core of its BTB/POZ domain and the oxidation of C-terminal cysteines. Plant Cell 18, 3670–3685 (2006).

23. Wang, D., Amornsiripanitch, N. & Dong, X. A genomic approach to identify regulatory nodes in the transcriptional network of systemic acquired resistance in plants. PLoS Pathog 2, e123 (2006).

24. Aarts, N. et al. Different requirements for EDS1 and NDR1 by disease resistance genes define at least two R gene-mediated signaling pathways in Arabidopsis. Proc Natl Acad Sci U S A 95, 10306–10311 (1998).

25. Falk, A. et al. EDS1, an essential component of R gene-mediated disease resistance in Arabidopsis has homology to eukaryotic lipases. Proc Natl Acad Sci U S A 96, 3292–3297 (1999).

26. Bhattacharjee, S., Halane, M.K., Kim, S.H. & Gassmann, W. Pathogen effectors target Arabidopsis EDS1 and alter its interactions with immune regulators. Science 334, 1405–1408 (2011).

27. Heidrich, K. et al. Arabidopsis EDS1 connects pathogen effector recognition to cell compartment-specific immune responses. Science 334, 1401–1404 (2011).

28. Rietz, S. et al. Different roles of Enhanced Disease Susceptibility1 (EDS1) bound to and dissociated from Phytoalexin Deficient4 (PAD4) in Arabidopsis immunity. New Phytol 191, 107–119 (2011).

29. Wagner, S. et al. Structural basis for signaling by exclusive EDS1 heteromeric complexes with SAG101 or PAD4 in plant innate immunity. Cell Host Microbe 14, 619–630 (2013).

30. Lapin, D. et al. A Coevolved EDS1-SAG101-NRG1 Module Mediates Cell Death Signaling by TIR-Domain Immune Receptors. Plant Cell 31, 2430–2455 (2019).

31. Bartsch, M. et al. Salicylic acid-independent ENHANCED DISEASE SUSCEPTIBILITY1 signaling in Arabidopsis immunity and cell death is regulated by the monooxygenase FMO1 and the Nudix hydrolase NUDT7. Plant Cell 18, 1038–1051 (2006).

32. Lapin, D., Bhandari, D.D. & Parker, J.E. Origins and Immunity Networking Functions of EDS1 Family Proteins. Annu Rev Phytopathol (2020).

33. Pajerowska-Mukhtar, K.M., Emerine, D.K. & Mukhtar, M.S. Tell me more: roles of NPRs in plant immunity. Trends Plant Sci 18, 402–411 (2013).

34. Chen, H. et al. A Bacterial Type III Effector Targets the Master Regulator of Salicylic Acid Signaling, NPR1, to Subvert Plant Immunity. Cell Host Microbe 22, 777–788 e777 (2017).

35. Sawyer, I.A. & Dundr, M. Nuclear bodies: Built to boost. J Cell Biol 213, 509–511 (2016).

36. Mosavi, L.K., Cammett, T.J., Desrosiers, D.C. & Peng, Z.Y. The ankyrin repeat as molecular architecture for protein recognition. Protein Science 13, 1435–1448 (2004).

37. Aravind, L. & Koonin, E.V. Fold prediction and evolutionary analysis of the POZ domain: Structural and evolutionary relationship with the potassium channel tetramerization domain. Journal of Molecular Biology 285, 1353–1361 (1999).

38. Cao, H., Bowling, S.A., Gordon, A.S. & Dong, X. Characterization of an Arabidopsis Mutant That Is Nonresponsive to Inducers of Systemic Acquired Resistance. Plant Cell 6, 1583–1592 (1994).

39. Delaney, T.P., Friedrich, L. & Ryals, J.A. Arabidopsis signal transduction mutant defective in chemically and biologically induced disease resistance. Proc Natl Acad Sci U S A 92, 6602–6606 (1995).

40. Glazebrook, J., Rogers, E.E. & Ausubel, F.M. Isolation of Arabidopsis mutants with enhanced disease susceptibility by direct screening. Genetics 143, 973–982 (1996).

41. Shah, J., Tsui, F. & Klessig, D.F. Characterization of a salicylic acid-insensitive mutant (sai1) of Arabidopsis thaliana, identified in a selective screen utilizing the SA-inducible expression of the tms2 gene. Molecular Plant-Microbe Interactions 10, 69–78 (1997).

42. Wang, W. et al. Structural basis of salicylic acid perception by Arabidopsis NPR proteins. Nature (2020).

43. Nishimura, M.T., Monteiro, F. & Dangl, J.L. Treasure your exceptions: unusual domains in immune receptors reveal host virulence targets. Cell 161, 957–960 (2015).

44. Bent, A.F. et al. RPS2 of Arabidopsis thaliana: a leucine-rich repeat class of plant disease resistance genes. Science 265, 1856–1860 (1994).

45. Garcia, A.V. et al. Balanced nuclear and cytoplasmic activities of EDS1 are required for a complete plant innate immune response. PLoS Pathog 6, e1000970 (2010).

46. Feys, B.J., Moisan, L.J., Newman, M.A. & Parker, J.E. Direct interaction between the Arabidopsis disease resistance signaling proteins, EDS1 and PAD4. Embo Journal 20, 5400–5411 (2001).

47. van Loon, L.C., Rep, M. & Pieterse, C.M. Significance of inducible defense-related proteins in infected plants. Annu Rev Phytopathol 44, 135–162 (2006).

48. Uknes, S. et al. Acquired resistance in Arabidopsis. Plant Cell 4, 645–656 (1992).

49. Zhou, J.M. et al. NPR1 differentially interacts with members of the TGA/OBF family of transcription factors that bind an element of the PR-1 gene required for induction by salicylic acid. Mol Plant Microbe Interact 13, 191–202 (2000).

50. Shearer, H.L. et al. Arabidopsis clade I TGA transcription factors regulate plant defenses in an NPR1-independent fashion. Mol Plant Microbe Interact 25, 1459–1468 (2012).

51. Zhang, Y., Tessaro, M.J., Lassner, M. & Li, X. Knockout analysis of Arabidopsis transcription factors TGA2, TGA5, and TGA6 reveals their redundant and essential roles in systemic acquired resistance. Plant Cell 15, 2647–2653 (2003).

52. Zhou, M. et al. WRKY70 prevents axenic activation of plant immunity by direct repression of SARD1. New Phytol 217, 700–712 (2018).

53. Xu, X.P., Chen, C.H., Fan, B.F. & Chen, Z.X. Physical and functional interactions between pathogen-induced Arabidopsis WRKY18, WRKY40, and WRKY60 transcription factors. Plant Cell 18, 1310–1326 (2006).

54. Eulgem, T. & Somssich, I.E. Networks of WRKY transcription factors in defense signaling. Curr Opin Plant Biol 10, 366–371 (2007).

55. Spoel, S.H. et al. Proteasome-mediated turnover of the transcription coactivator NPR1 plays dual roles in regulating plant immunity. Cell 137, 860–872 (2009).

56. Bharti, K. et al. Isolation and characterization of HsfA3, a new heat stress transcription factor of Lycopersicon peruvianum. Plant J 22, 355–365 (2000).

57. Chen, H. et al. Arabidopsis DREB2C functions as a transcriptional activator of HsfA3 during the heat stress response. Biochem Biophys Res Commun 401, 238–244 (2010).

58. Ma, J. & Ptashne, M. A new class of yeast transcriptional activators. Cell 51, 113–119 (1987).

59. Hope, I.A., Mahadevan, S. & Struhl, K. Structural and functional characterization of the short acidic transcriptional activation region of yeast GCN4 protein. Nature 333, 635–640 (1988).

60. Pennica, D. et al. The amino acid sequence of murine p53 determined from a c-DNA clone. Virology 134, 477–482 (1984).

61. Cress, W.D. & Triezenberg, S.J. Critical structural elements of the VP16 transcriptional activation domain. Science 251, 87–90 (1991).

62. Conaway, R.C. & Conaway, J.W. Function and regulation of the Mediator complex. Curr Opin Genet Dev 21, 225–230 (2011).

63. Galbraith, M.D., Donner, A.J. & Espinosa, J.M. CDK8: a positive regulator of transcription. Transcription 1, 4–12 (2010).

64. Zhu, Y. et al. CYCLIN-DEPENDENT KINASE8 differentially regulates plant immunity to fungal pathogens through kinase-dependent and -independent functions in Arabidopsis. Plant Cell 26, 4149–4170 (2014).

65. Chen, J. et al. NPR1 Promotes Its Own and Target Gene Expression in Plant Defense by Recruiting CDK8. Plant Physiol 181, 289–304 (2019).

66. Jin, H. et al. Salicylic acid-induced transcriptional reprogramming by the HAC-NPR1-TGA histone acetyltransferase complex in Arabidopsis. Nucleic Acids Res 46, 11712–11725 (2018).

67. Lebel, E. et al. Functional analysis of regulatory sequences controlling PR-1 gene expression in Arabidopsis. Plant J 16, 223–233 (1998).

68. Kesarwani, M., Yoo, J.M. & Dong, X.N. Genetic interactions of TGA transcription factors in the regulation of pathogenesis-related genes and disease resistance in Arabidopsis. Plant Physiology 144, 336–346 (2007).

69. Liu, L. et al. Salicylic acid receptors activate jasmonic acid signalling through a non-canonical pathway to promote effector-triggered immunity. Nat Commun 7, 13099 (2016).

70. Chang, M. et al. PBS3 Protects EDS1 from Proteasome-Mediated Degradation in Plant Immunity. Mol Plant 12, 678–688 (2019).

71. Century, K.S., Holub, E.B. & Staskawicz, B.J. NDR1, a locus of Arabidopsis thaliana that is required for disease resistance to both a bacterial and a fungal pathogen. Proc Natl Acad Sci U S A 92, 6597–6601 (1995).

72. Warren, R.F., Merritt, P.M., Holub, E. & Innes, R.W. Identification of three putative signal transduction genes involved in R gene-specified disease resistance in Arabidopsis. Genetics 152, 401–412 (1999).

73. Nawrath, C., Heck, S., Parinthawong, N. & Metraux, J.P. EDS5, an essential component of salicylic acid-dependent signaling for disease resistance in Arabidopsis, is a member of the MATE transporter family. Plant Cell 14, 275–286 (2002).

74. Zheng, Z., Qualley, A., Fan, B., Dudareva, N. & Chen, Z. An important role of a BAHD acyl transferase-like protein in plant innate immunity. Plant J 57, 1040–1053 (2009).

75. Ryals, J. et al. The Arabidopsis NIM1 protein shows homology to the mammalian transcription factor inhibitor I kappa B. Plant Cell 9, 425–439 (1997).

76. Feys, B.J. et al. Arabidopsis SENESCENCE-ASSOCIATED GENE101 stabilizes and signals within an ENHANCED DISEASE SUSCEPTIBILITY1 complex in plant innate immunity. Plant Cell 17, 2601–2613 (2005).

77. Saleh, A. et al. Posttranslational Modifications of the Master Transcriptional Regulator NPR1 Enable Dynamic but Tight Control of Plant Immune Responses. Cell Host Microbe 18, 169–182 (2015).

78. Cho, W.K. et al. Mediator and RNA polymerase II clusters associate in transcription-dependent condensates. Science 361, 412–415 (2018).

79. Plys, A.J. & Kingston, R.E. Dynamic condensates activate transcription. Science 361, 329–330 (2018).

80. Zavaliev, R., Mohan, R., Chen, T. & Dong, X. Formation of NPR1 Condensates Promotes Cell Survival during the Plant Immune Response. Cell 182, 1093–1108 e1018 (2020).

81. Venugopal, S.C. et al. Enhanced disease susceptibility 1 and salicylic acid act redundantly to regulate resistance gene-mediated signaling. PLoS Genet 5, e1000545 (2009).

82. DeFraia, C.T., Zhang, X.D. & Mou, Z.L. Elongator subunit 2 is an accelerator of immune responses in Arabidopsis thaliana. Plant Journal 64, 511–523 (2010).

83. Cui, H. et al. A core function of EDS1 with PAD4 is to protect the salicylic acid defense sector in Arabidopsis immunity. New Phytol 213, 1802–1817 (2017).

84. Cui, H. et al. Antagonism of Transcription Factor MYC2 by EDS1/PAD4 Complexes Bolsters Salicylic Acid Defense in Arabidopsis Effector-Triggered Immunity. Mol Plant 11, 1053–1066 (2018).

85. Rusterucci, C., Aviv, D.H., Holt, B.F., 3rd, Dangl, J.L. & Parker, J.E. The disease resistance signaling components EDS1 and PAD4 are essential regulators of the cell death pathway controlled by LSD1 in Arabidopsis. Plant Cell 13, 2211–2224 (2001).

86. Thaler, J.S., Humphrey, P.T. & Whiteman, N.K. Evolution of jasmonate and salicylate signal crosstalk. Trends Plant Sci 17, 260–270 (2012).

87. Spoel, S.H. et al. NPR1 modulates cross-talk between salicylate- and jasmonate-dependent defense pathways through a novel function in the cytosol. Plant Cell 15, 760–770 (2003).

88. Wirthmueller, L., Zhang, Y., Jones, J.D. & Parker, J.E. Nuclear accumulation of the Arabidopsis immune receptor RPS4 is necessary for triggering EDS1-dependent defense. Curr Biol 17, 2023–2029 (2007).

89. Zhang, X., Wang, C., Zhang, Y., Sun, Y. & Mou, Z. The Arabidopsis mediator complex subunit16 positively regulates salicylate-mediated systemic acquired resistance and jasmonate/ethylene-induced defense pathways. Plant Cell 24, 4294–4309 (2012).

90. Zhang, Y. et al. A Highly Efficient Agrobacterium-Mediated Method for Transient Gene Expression and Functional Studies in Multiple Plant Species. Plant Commun 1, 100028 (2020).

91. Chen, H. et al. Firefly luciferase complementation imaging assay for protein-protein interactions in plants. Plant Physiol 146, 368–376 (2008).

92. Saleh, A., Alvarez-Venegas, R. & Avramova, Z. An efficient chromatin immunoprecipitation (ChIP) protocol for studying histone modifications in Arabidopsis plants. Nat Protoc 3, 1018–1025 (2008).

