## Supplementary Figs. 1-6 and Tables 1-2 for "Two interacting transcriptional coactivators cooperatively control plant immune responses"

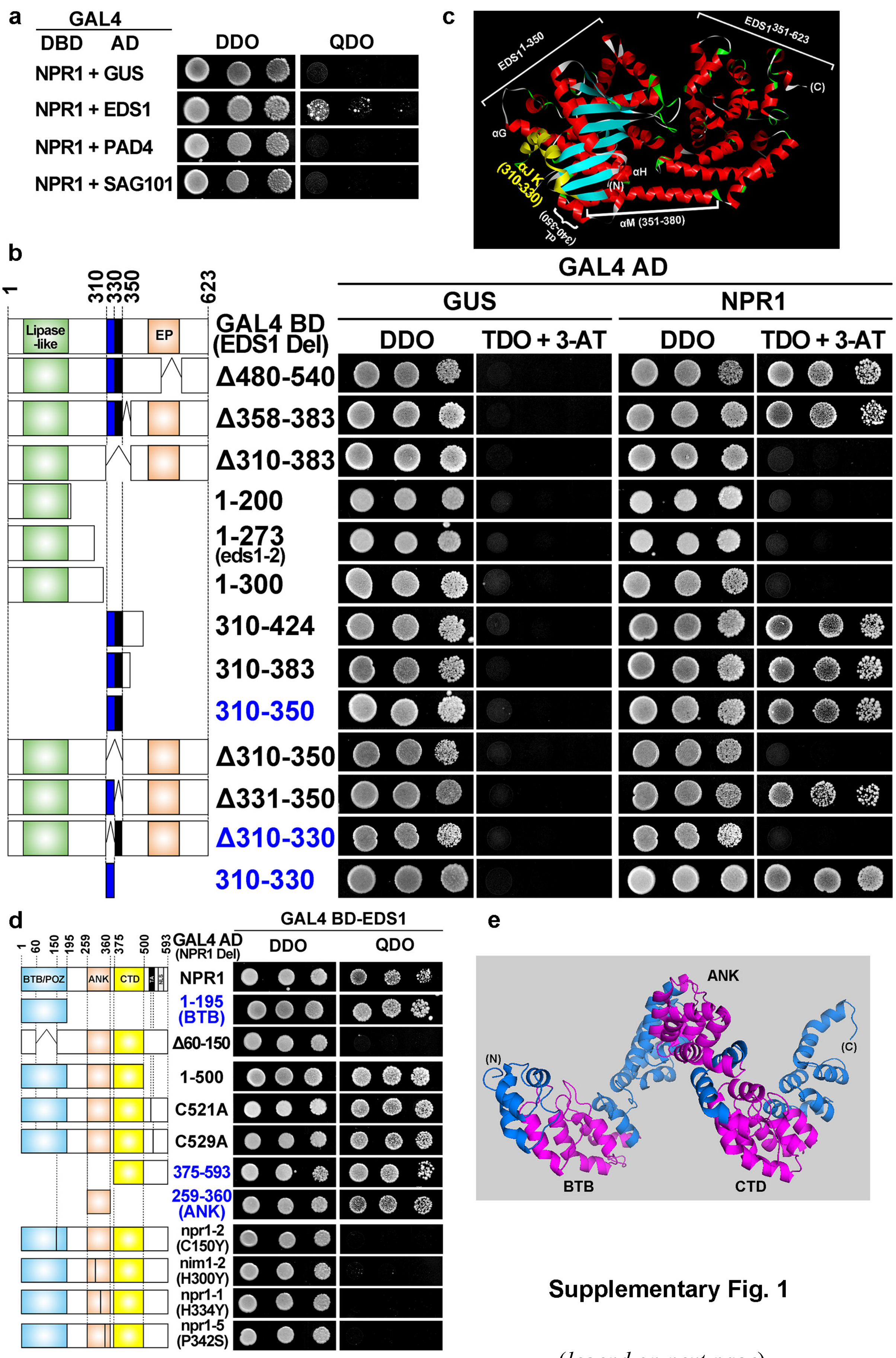


**Supplementary Fig. 1**

*(legend on next page)*

**Supplementary Fig. 1 | Identification of the interaction domains in NPR1 and EDS1. a**, NPR1 specifically interacts with EDS1. The yeast strain Y187 with full-length NPR1 fused with GAL4-DBD was mated with yeast strain AH109 harboring GAL4-AD fusion protein (GUS, PAD4 or SAG101), and then the diploid yeast cells were grown on synthetic double dropout agar media (DDO, without Leu and Trp) and selective quadruple dropout medium (QDO, without Leu, Trp, His, and Ade). GAL4-DBD, GAL4-DNA binding domain; AD, activation domain. **b**, NPR1 recognizes the α-helix region (310-330) in EDS1. Left panel is schematic diagram of the EDS1 deletion (Del) fragments. The numbers indicate the corresponding amino acid residues. The specific regions in EDS1, such as a lipase-like domain, a region (480-540) partially overlapping with EP (EDS1 and PAD4-defined) domain, a coiled-coil domain (358-383) defined by Pfam and MARCOIL, a region (310-350) encompassing two α-helices defined by Phyre$^2$, the region (1-273) that is almost equivalent to the left-over region in the null mutant *eds1-2*, a fast neutron-derived mutant containing truncated product S276 (serine 276)-Stop[25], and others, are shown in the diagram. Yeast cells were grown on DDO and triple dropout medium (TDO, without Leu, Trp and His) plus 1 mM 3-aminotriazole (3-AT). Diagram is drawn to scale. **c**, Overview of EDS1 protein 3D structure (PDB: 4NFU). The protein ribbon model with diverse structures is illustrated by Accelrys Discovery Studio Visualizer. The protruding interaction α helices JK (310-330) for NPR1-EDS1 interaction is highlighted in yellow. Other structures such as αG, αH, αL and αM are marked in white. Detail information of α helices are described as previously[29]. **d,e**, EDS1 dynamically interacts with NPR1. The left panel shown is a schematic diagram of the NPR1 deletion (Del) mutants. The numbers indicate the corresponding amino acid residues. BTB/POZ, Broad complex, Tramtrack, and Bric-a-brac/Pox virus and Zinc finger domain; ANK, ankyrin repeats; CTD, a C-terminal domain; TA, transactivation domain; C521A/C529A, the Ala substitution of two residues (Cys 521 and Cys 529) that are critical for TA; NLS, nuclear localization signal. Note that the N-terminal BTB/POZ domain and a second repression region (463-513) in CTD confer the transcriptional repression activity of NPR1 upon SA induction and under normal conditions, respectively[22]. Diagram is drawn to scale. Modelled structure of NPR1 was generated by Phyre$^2$ server using the template c4cj9A (**e**). PyMol was used to visualize and arrange modelled structure.

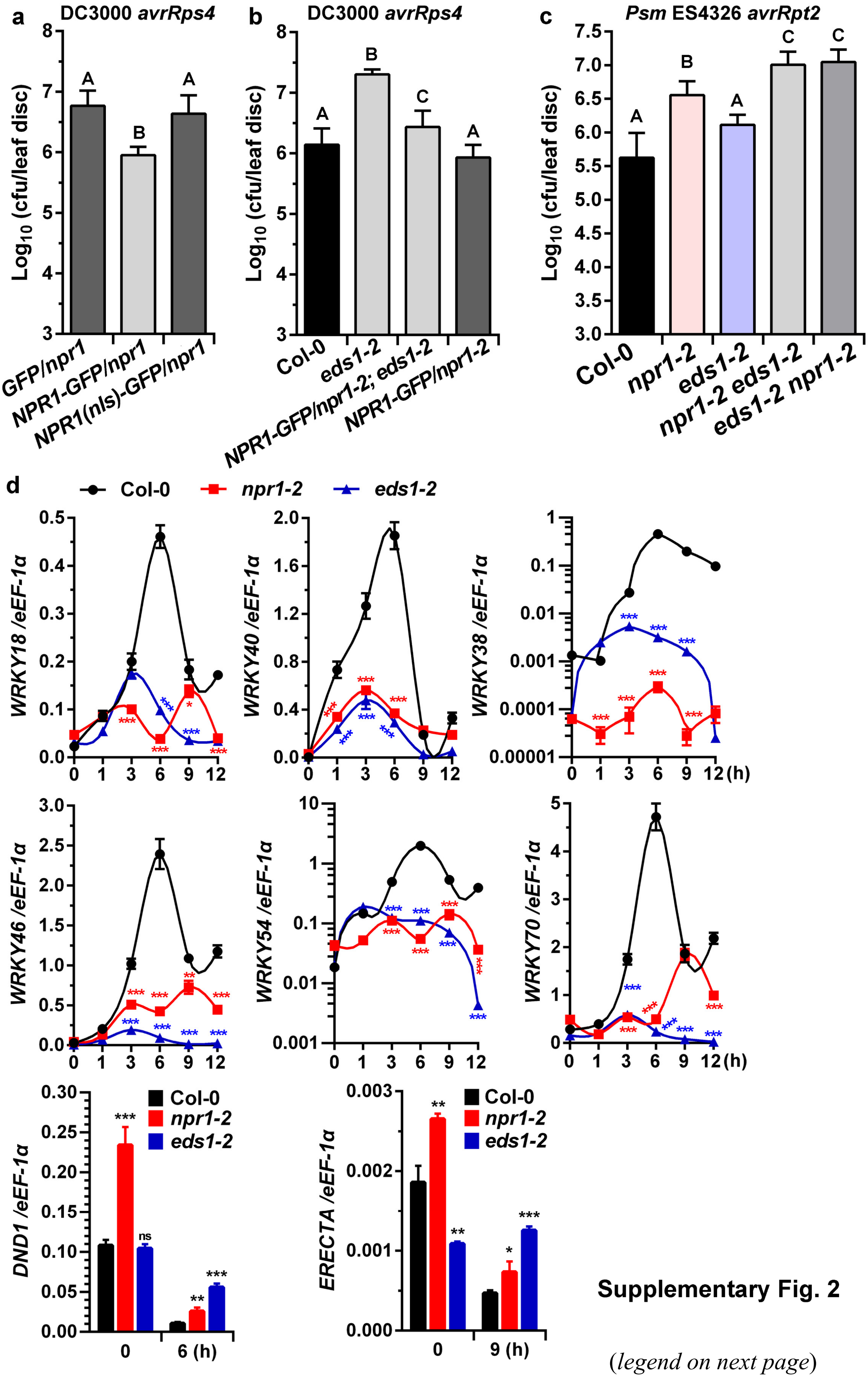


**Supplementary Fig. 2**

(*legend on next page*)

**Supplementary Fig. 2 | Genetic and molecular interactions between NPR1 and EDS1 in ETI. a**, Nuclear NPR1 is involved in ETI. Bacterial growth assays were performed using transgenic *Arabidopsis* constitutively expressing *GFP*, *NPR1-GFP* or *NPR1 (nls)-GFP* under the control of *CaMV* 35S promoter in the *npr1-2* background. **b**, NPR1 functions as a partner of EDS1 in TIR-NB-LRR R protein (RRS1S/RPS4)-activated ETI. Growth of bacterial was determined in Col-0, *eds1-2*, *35S:NPR1/npr1-2;eds1-2* and *35S:NPR1-GFP/npr1-2* transgenic plants. **c**, EDS1 functions as a partner of NPR1 in CC-NLR protein (RPS2)-activated ETI. Leaves of soil-grown plant were inoculated with *Pst* DC3000 *avrRps4* ($OD_{600}$ = 0.0005) in (**a**) and (**b**) or *Psm* ES4326 *avrRpt2* ($OD_{600}$ = 0.0002) in (**c**); the bacterial titers were measured at 2 dpi (CFU, colony-forming units). Error bars represent standard deviation (SD); n = 6 biologically independent samples (**a-c**). Statistically significant difference is indicated by different lowercase letters ($P < 0.01$). **d**, NPR1 and EDS1 co-regulate a set of common target genes in RRS1S/RPS4-activated ETI. EDS1-induced *WRKY* genes (*WRKY18*, *WRKY40*, *WRKY38*, *WRKY46*, *WRKY54* and *WRKY70*) and EDS1-repressed genes (*DND1* and *ERECTA*) were analyzed using real-time qPCR. Leaves from 4-week-old soil-grown plants were inoculated with the avirulent bacterial pathogen *Pst* DC3000 *avrRps4* ($OD_{600}$ = 0.01). Total RNA was extracted from inoculated leaves collected at the indicated time points. Expression of *WRKY38* and *WRKY54* was plotted on a $\log_{10}$ scale; gene expression levels were normalized against the constitutively expressed *eEF-1α*. Error bars indicate the (±) SD (n = 3 biologically independent samples). Statistical difference is shown between Col-0 and single mutant (*npr1-2* or *eds1-2*) plants (*t*-test, *, $P < 0.05$; **, $P < 0.01$; ***, $P < 0.001$). ns, no significant difference.

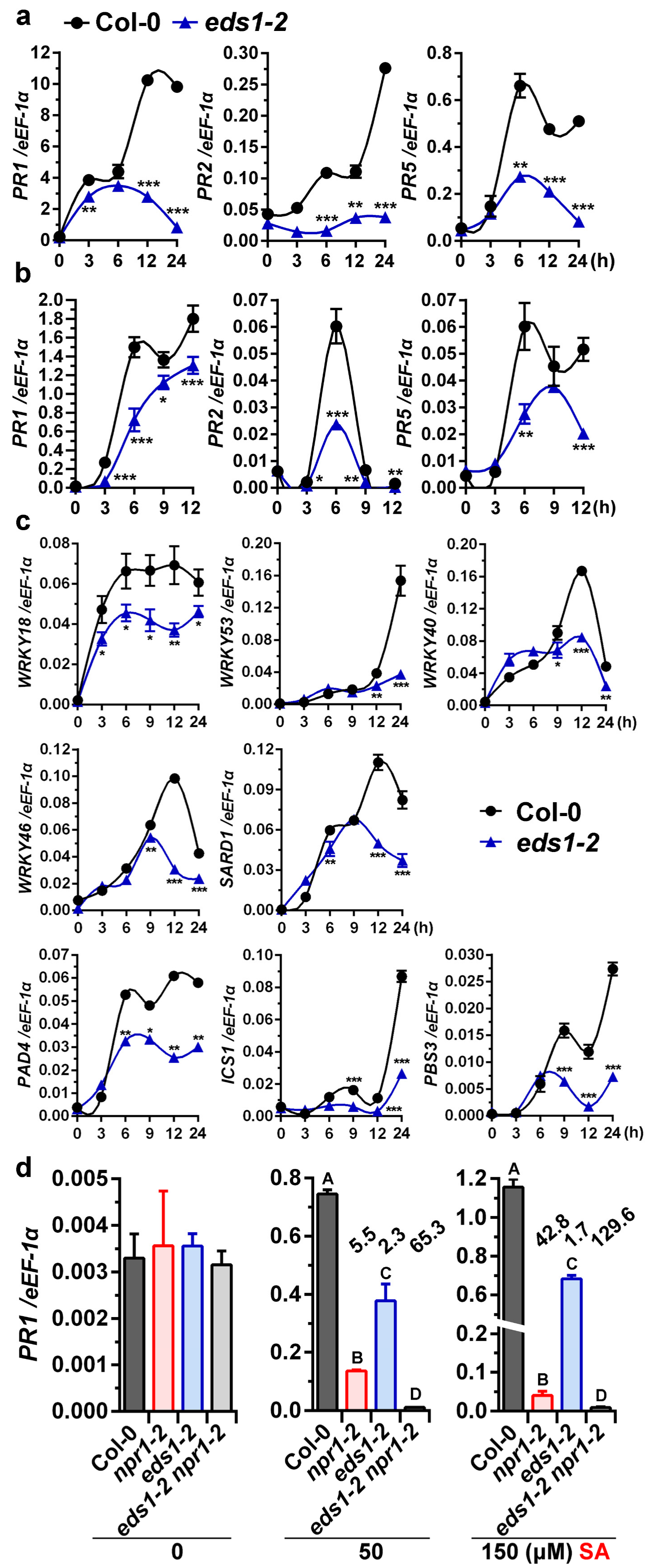


**Supplementary Fig. 3**

(*legend on next page*)

**Supplementary Fig. 3│EDS1 and NPR1 coregulate some SA-responsive genes. a,b**, EDS1 can functions as a positive regulator of SA signaling (downstream of SA). Total RNA was extracted from rosette leaves of 4-week-old soil-grown plants (**a**) infiltrated with 0.3 mM SA or from 2-week-old seedlings grown on one-half strength Murashige and Skoog (1/2 MS) solid agar media (**b**) exogenously treated with hydroponic 0.5 mM SA solution. Expression of *PR1*, *PR2*, and *PR5* in Col-0 and *eds1-2* was analyzed using real-time qPCR. **c**, EDS1 upregulates some NPR1-mediated and SA-responsive genes. Total RNA was extracted from 2-week-old seedlings on 1/2 MS media exogenously treated with hydroponic 0.5 mM SA solution. Expression of a subset of *WRKY* genes (*WRKY18*, *WRKY40*, *WRKY46*, and *WRKY53*), SA accumulation (*SARD1* and *PAD4*) and biosynthesis (*ICS1* and *PBS3*) genes were analyzed using qPCR. Expression was normalized against constitutively expressed *eEF-1α*. Error bars represent ± SD; n = 3 biologically independent samples (**a-c**). Statistical significance comparisons between Col-0 and *eds1-2* for each time point were shown on the bars (*t*-test, *, $P < 0.05$; **, $P < 0.01$; ***, $P < 0.001$). **d**, NPR1 and EDS1 synergistically activate *PR1* in response to SA. Total RNA was isolated from 2-week-old seedlings grown on 1/2 MS agar plates containing different low concentrations of SA (50 and 150 μM) for long-term treatment (mimic SAR treatment). Error bars represent SD (n = 4 biologically independent samples). Different letters indicate statistical difference (two-way ANOVA, $P < 0.01$). The folds above the error bars indicate the fold change for the gene expression reduction compared with the value obtained in Col-0.

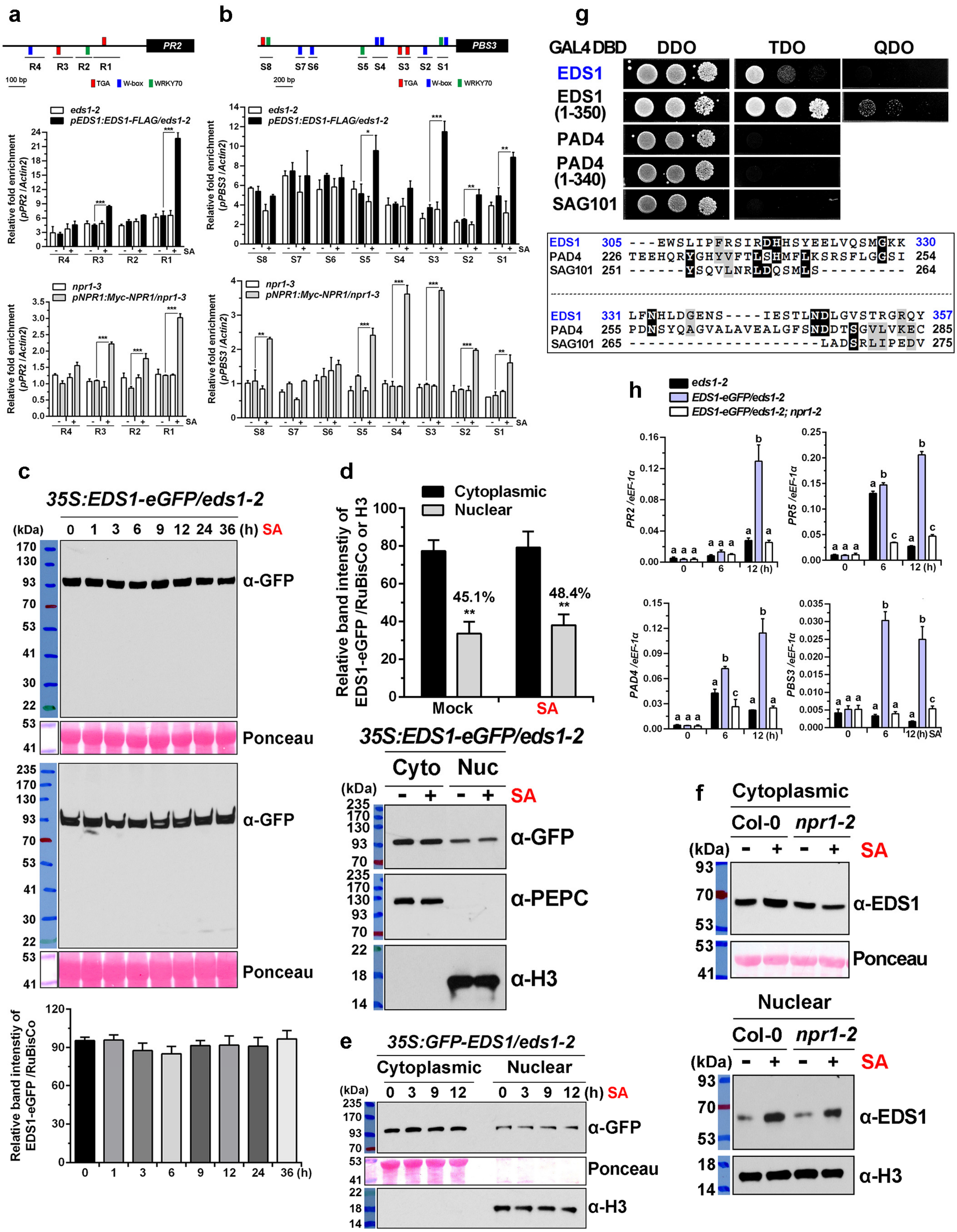


**Supplementary Fig. 4**

*(legend on next page)*

(legend on next page)

**Supplementary Fig. 4 | Recruitment of NPR1-EDS1 complex onto promoters of plant defense genes upon SA induction. a,b**, The NPR1-EDS1 protein complex occupies specific regions of the *PR2* and *PBS3* promoters. ChIP-qPCR shows enrichment of NPR1 and EDS1 at the *PR2* (**a**) and *PBS3* (**b**) genomic loci. The mutant control and transgenic plants were harvested after 0.5 mM SA (+) or water (-) treatment for 9 h. After sonication and clearance, ChIP assays were performed with anti-FLAG magnetic beads or anti-Myc antibody. The amount of immunoprecipitated DNA was quantified by qPCR using primers specific to each amplicon. Samples from *npr1-2* or *eds1-2* mutants were used as negative controls. Specific enrichment of fusion protein on genomic DNA sequences was calculated relative to that on *Actin2* coding DNA sequences. Error bars indicate SD (n =3 biologically independent samples). Significances of differences are shown (*t*-test, *, $P < 0.05$; **, $P < 0.01$; ***, $P < 0.001$). Top panel is the schematic representation of the *cis*-elements and chromatin fragments (R1 to R4 for *PR2* and S1 to S8 for *PBS3*) of amplicon in the genomic regions. Diagram is drawn to scale. Details on the positions of the primers are provided in Supplementary Table 1. W-box (blue box), TTGACT/C; TGA motif (red box), TGACG; TGA2/W (chimeric color), TGACTT (TGA2 binding sites overlapping W-box); WRKY70 (green box), GACTTTT (putative WRKY70 binding sites)[52]. The perfect *cis*-elements with inverted consensus sequences are shown below the promoter. **c**, SA does not induce constitutively expressed EDS1-eGFP. Transgenic plants constitutively expressing EDS1-eGFP under the control of the *CaMV* 35S promoter in the *eds1-2* background (*35S*:*EDS1-eGFP*/*eds1-2*) were treated with 0.5 mM SA solution and collected at the indicated time points. Total protein was analyzed in western blotting using anti-GFP antibody. The ribulose-1,5-bisphosphate carboxylase/oxygenase (RuBisCo) large subunit bands visualized by Ponceau S staining are shown as protein loading controls. Lower panel shows relative EDS1-eGFP band intensity of immunoblot analysis. Blots were quantitated using the ImageJ software. Protein level was normalized to the constitutively expressed RuBisCo. Bars indicate SE from three independent blots. **d,e**, SA does not facilitate nuclear translocation of EDS1. The *35S*:*EDS1-eGFP*/*eds1-2* transgenic plants (**d**) and *35S*:*GFP-EDS1*/*eds1-2* plants (**e**) were treated with 0.5 mM SA solution. The former plants were harvested 9 h after treatment for cell fractionation; the latter were collected at the indicated time points. EDS1-eGFP and GFP-EDS1 proteins were immunologically detected using an anti-GFP antibody. Anti-phosphoenolpyruvate carboxylase (PEPC) and Ponceau S staining were used for detection of cytoplasmic (Cyto) markers; anti-Histone H3 antibody was used for detection of the nuclear (Nuc) marker. Upper panel, relative EDS1-eGFP band intensity of immunoblot analysis. Blots were quantitated using the ImageJ software. Cytoplasmic protein level was normalized to the constitutively expressed RuBisCo; nuclear protein level was normalized to histone H3. Bars indicate SE from four independent blots. Significances of differences between cytoplasmic and nuclear fractions are analyzed for each treatment (*t*-test, **, $P < 0.01$). **f**, SA induces the accumulation of endogenous EDS1 in cytoplasm and nuclei. Nucleo-cytoplasmic distribution of endogenous EDS1 protein in SA-treated plant cells was performed using 5-week-old soil-grown plants that were harvested after 12 h treatment with foliar sprays of 0.5 mM SA solution. EDS1 and histone H3 proteins were immunologically detected using anti-EDS1 and anti-histone H3 antibodies, respectively. **g,** EDS1 specifically possesses autoactivation, but not PAD4 or SAG101. The yeast strain Y187 with indicated protein fused with GAL4-DBD was mated with yeast strain AH109 harboring GUS fused with GAL4-AD. Diploid yeast cells were grown on synthetic dropout agar media such as DDO (double dropout; without Leu, Trp), TDO (triple dropout; without Leu, Trp, His) and QDO (quadruple dropout; without Leu, Trp, His, and Ade). Bottom, multiple sequence alignment of NPR1-interacting domain (310~330) and TAD (331~350) of EDS1 among EDS1 and its partners using ClustalW and BoxShade servers (SIB Swiss Institute of Bioinformatics). The numbers indicate the corresponding amino acid residues. **h**, SA-induced

EDS1 activates plant defense genes. Expression of *PR2*, *PR5*, *PAD4* and *PBS3* was analyzed by real-time qPCR. Expression was normalized against constitutively expressed *eEF-1α*. Four-week-old soil-grown plants were infiltrated with 0.5 mM SA solution and leaf tissues were collected at indicated time points. Bars represent SD; n = 4 biologically independent samples. Different letters indicate significant differences (Two-way ANOVA, Tukey's multiple comparisons test, $P < 0.05$). The statistical comparisons were made separately among *eds1-2*, *35S:EDS1-eGFP/eds1-2* and *35S:EDS1-eGFP/eds1-2;npr1-2* for each time point.

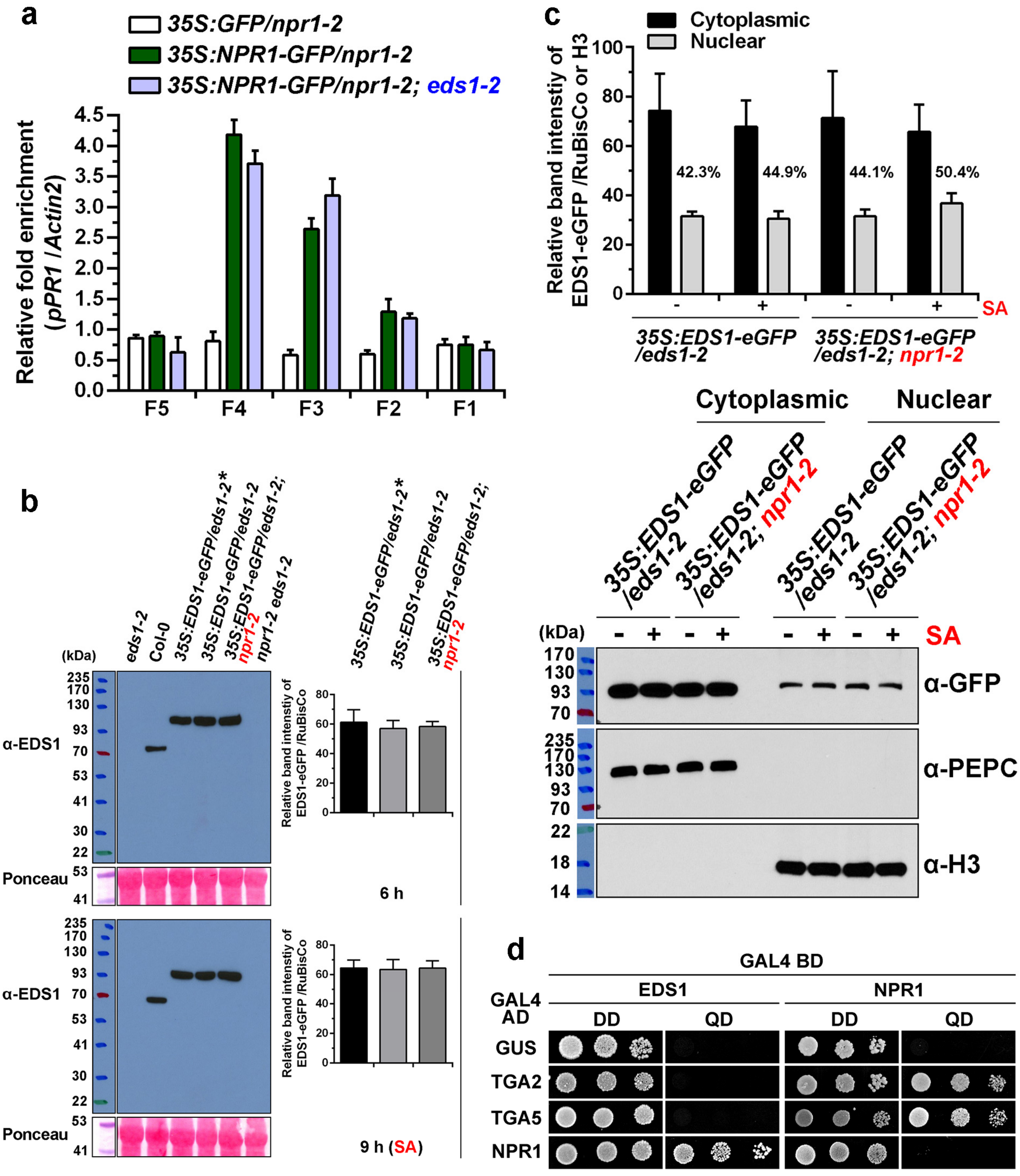


**Supplementary Fig. 5**

(*legend on next page*)

**Supplementary Fig. 5 | A physical NPR1-EDS1 interaction confers the recruitment of EDS1 to *PR1* promoter. a**, EDS1 slightly influences the association of NPR1-GFP with the *PR1* promoter. Four-week-old soil-grown transgenic plants were treated with foliar sprays plus soil drenches of 0.5 mM SA solution for 9 h. The collected leaf tissues were subjected to ChIP assay using GFP-Trap magnetic beads. Schematic representation of the chromatin fragments (F1 to F5) is illustrated in Fig. 4a. Bars indicate SD (n = 3 biological independent samples). This experiment was repeated twice with similar results. **b**, NPR1 has no significant effect on the expression of EDS1-eGFP. Total protein prepared from indicated genotypes was analyzed by immunoblotting using an anti-EDS1 antibody. Relative band intensity was analyzed by normalizing EDS1-eGFP protein against the corresponding RuBisCo. Error bars indicate SD from triplicate technical repeats. The *35S*:*EDS1-eGFP*/*eds1-2* denoted by asterisk (*) is the original homozygous transgenic line that was used to cross with *npr1-2*; the resulting homozygous progenies (i.e. *35S*:*EDS1-eGFP*/*eds1-2* and *35S*:*EDS1-eGFP*/*eds1-2*; *npr1-2*) from same generation were identified. **c**, NPR1 does not facilitate nuclear translocation of EDS-eGFP. The indicated transgenic seedlings were treated with 0.5 mM SA (+) or water (-) for 9 h. Relative band intensity and immunoblotting are performed as described in Supplementary Fig. 4d-f. Percentage indicates the ratio of nuclear to cytoplasmic distribution. Bars indicate SE from four independent experiments. **d**, The interaction between EDS1 and TGA2/5 is not detected in Y2H assays. Yeast cells were grown on synthetic double dropout agar media (DD, without Leu and Trp) and selective quadruple dropout medium (QD, without Leu, Trp, His, and Ade). GAL4 BD, GAL4-DNA binding domain; AD, activation domain.

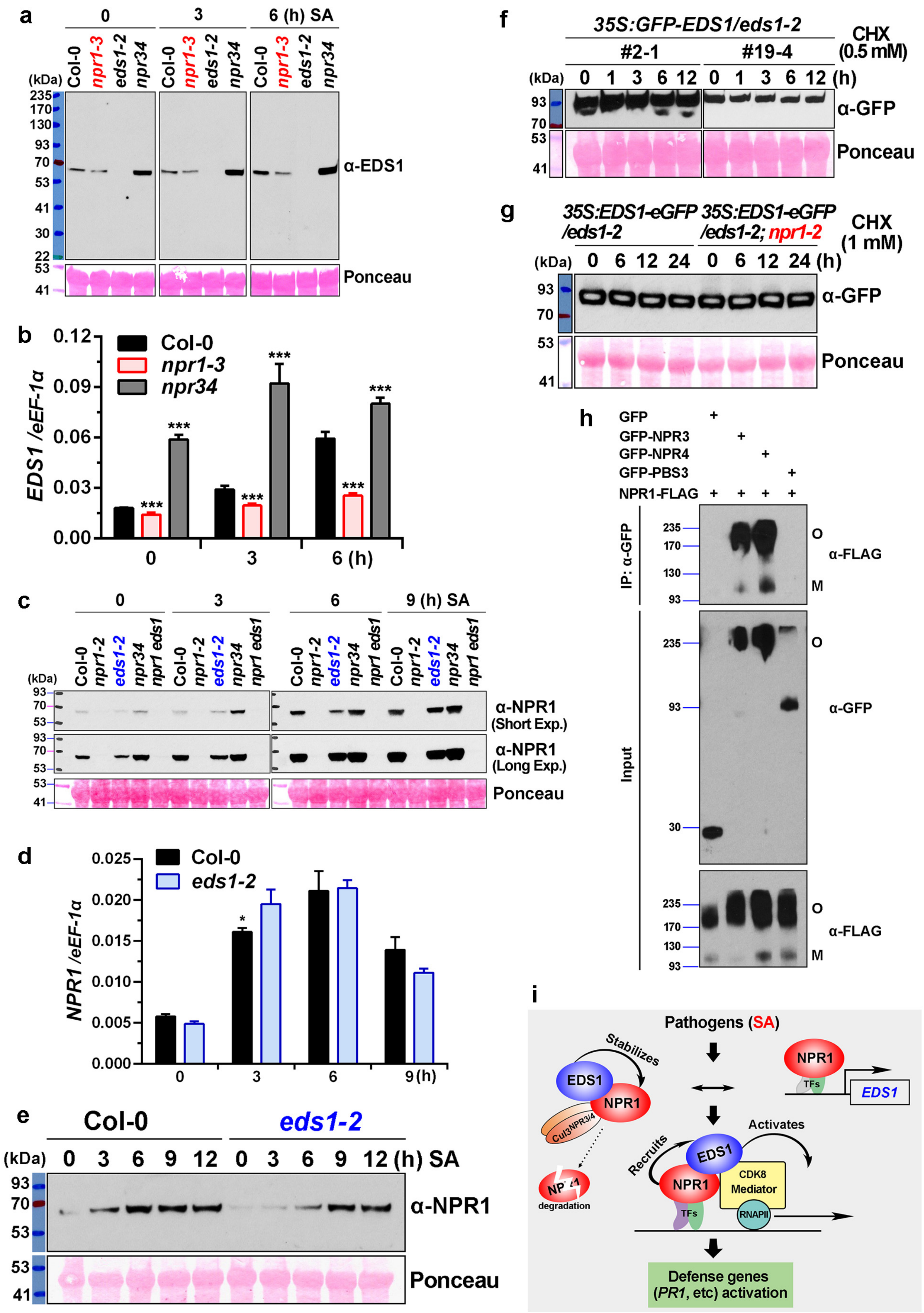


**Supplementary Fig. 6**

(*legend on next page*)

**Supplementary Fig. 6 | NPR1 transcriptionally upregulates *EDS1* expression, while EDS1 stabilizes NPR1 protein.** **a,b**, NPR1 transcriptionally upregulates *EDS1* expression. EDS1 protein level (**a**) and EDS1 transcription level (**b**) are analyzed in Col-0, *npr1-3*, *eds1-2* and *npr3-2 npr4-2* (*npr34*) seedlings exogenously treated with hydroponic 0.5 mM SA solution for indicated times. **c-e**, EDS1 stabilizes NPR1 in SA-mediated defense responses. NPR1 protein level (**c**) and *NPR1* transcription level (**d**) are investigated in Col-0, *npr1-2*, *eds1-2*, *npr3-2 npr4-2* (*npr34*), and *npr1-2 eds1-2* (*npr1 eds1*) seedlings treated with 0.5 mM SA solution for indicated times. The short exposure and long exposure (Exp.) of same protein gel blot are shown. Total RNA was extracted from 2-week-old 1/2 MS media-grown seedlings and subjected to real-time qPCR analysis (**b,d**). The *eEF-1α* gene was used as an internal control. **f,g**, EDS1 protein turnover is not affected by NPR1. The constitutively expressed EDS1 fusion proteins (GFP-EDS1 and EDS1-eGFP) in indicated transgenic seedlings are constant in the presence of cycloheximide (CHX) even at high concentrations (0.5 and 1 mM) and for a long duration (24 h). **h**, Oligomeric and monomeric NPR1 associates with NPR3/4 in plants. *N. benthamiana* was co-transformed with indicated constructs such as *35S*:*GFP*, *35S*:*GFP-NPR3*/*4*/*PBS3* and p*NPR1*:*NPR1-3xFLAG*. Approximately 0.2 g of leaf tissues was used to total protein extraction. Co-IP and immunoblotting assays were performed under reducing conditions. O, oligomeric GFP-NPR1/3/4; M, monomeric GFP-NPR1/3/4. **i**, A working model of EDS1-mediated functions in plant immunity. Under plant-pathogen interactions and SA induction, EDS1, acting as a transcriptional coactivator, is directed recruited by SA-induced NPR1 onto promoters of defense genes, and in turn EDS1 cooperates with NPR1 and directly interacts with Meditator complex (e.g. CDK8 module) to facilitate the assemble of RNAPII preinitiation complex, thereby potentiating transcription of downstream defense genes., In addition, EDS1 prevents NPR1 from degradation under normal and immune induced conditions, providing optimal NPR1 level to further immune responses. Simultaneously NPR1 directly targets *EDS1* and upregulates its transcription upon immune induction. Bars indicate (±) SD; n = 4 biological independent samples (**b,d**). Statistical significances of difference from control were shown for each time point (*t*-test, *, $P < 0.05$; **, $P < 0.01$; ***, $P < 0.001$). Total protein was analyzed by reducing SDS-PAGE and immunoblotting using indicated proteins. RuBisCo protein stained by Ponceau S was used as internal control. Protein sizes marked on the left are in kDa.

**Supplementary Table 1. Primers Used in This Study**

| Primer Name | Primer Sequence (5’ - 3’) |
|---|---|
| **Plasmid and Cloning** | |
| GW_F (attB1) | ggggACAAGTTTGTACAAAAAAGCAGGCTtc |
| GW_R (attB2) | gggcACCACTTTGTACAAGAAAGCTGGGTc |
| pNPR1:NPR1 | GW_F_TTTATGAATATTTAATCTGATTTTTTGGCT |
| | GW_R_ CCGACGACGATGAGAGAGTTTAC |
| pEDS1:EDS1 | GW_F_ GTTTATCAGATTCCACGTACGAT |
| | GW_R_ GGTATCTGTTATTTCATCCATCA |
| EDS1 | GW_F_ATGGCGTTTGAAGCTCTTACC |
| | GW_R_GGTATCTGTTATTTCATCCATCA |
| PAD4 | GW_F_ATGGACGATTGTCGATTCGAG |
| | GW_R_AGTCTCCATTGCGTCACTCTC |
| SAG101 | GW_F_ATGGAGTCTTCTTCTTCACTAAAAG |
| | GW_R_TTGTGACTTACCATAACTCTCGT |
| EDS1Δ480-540 | EDS1_F_ATGGCGTTTGAAGCTCTTACC |
| | EDS1_R_GGTATCTGTTATTTCATCCATCATATAGTC |
| | EDS1_R1437_AAGCATGATCCGCACTCGGATCCGT TCTTTAAATGTCGATGGTAGTTTGC |
| | EDS1_F1621_CTACCATCGACATTTAAAGAACGGAT CCGAGTGCGGATCAT |
| EDS1Δ358-383 | EDS1_R1071_ CTATCCATGCTAGTTTCTTTAAAAGT ACTGTCTGCCTCTTGTGCTC |
| | EDS1_F1150_AGTGAGCACAAGAGGCAGACAGTACT TTTTAAAGAAACTAGCATGGATAGAAG |
| EDS1Δ310-383 | EDS1_R930_ TTGGCTTGTATTCATCTTCTATCCA ATAGCTATGATGATCTCTGATACTTCGA |
| | EDS1_F1168_ AAGTATCAGAGATCATCATAGCTAT TGGATAGAAGATGAATACAAGCCA |
| EDS1Δ310-350 | EDS1_R930m3_CGTACTGTCTGCCTCTTGTGCT TGGAATTAGAGACCATTCTTGTTCATC |
| | EDS1_F1051_ AAGAATGGTCTCTAATTCCA AGCACAAGAGGCAGACAGT |
| EDS1Δ331-350 | EDS1_R990_ CGTACTGTCTGCCTCTTGTGCT CTTCTTTCCCATCGACTGTACC |
| | EDS1_F1051m5_ TGGTACAGTCGATGGGAAAGAAG AGCACAAGAGGCAGACAGT |
| EDS1Δ310-330 | EDS1_R930m4_ CTCCATCCAAATGATTAAACAA TGGAATTAGAGACCATTCTTGTTCATC |
| | EDS1_F991m4_ AAGAATGGTCTCTAATTCCA TTGTTTAATCATTTGGATGGAGAAAACT |

| | |
|---|---|
| EDS1_1-120 | GW_R_ TGTGAACACTATCTGTTTTCTACTCCT |
| EDS1_1-200 | GW_R_GACAAAGTTCACAAAGAACCG |
| EDS1_1-273 (eds1-2) | GW_R_CACTGCAACCAATCTCTTCTC |
| EDS1_1-300 | GW_R_GTGATCCTCTTTCAAAAGCAACTTG |
| EDS1_310-424 | GW_F310_ATGCCATTTCGAAGTATCAGAGATCATCATAG |
| | GW_R_ CTTGACGTTTGCTTTGAAGTCATTC |
| EDS1_310-383 | GW_R_ AAACCTCTCTTGCTCGATCAC |
| EDS1_310-350 | GW_R_ CACTCCAAGGTCATTGAGCGTAG |
| EDS1_310-330 | GW_R_ CTTCTTTCCCATCGACTGTACC |
| EDS1_357-623 | GW_F_ ATGTACGTTCAAGCTGCATTAGAGG |
| EDS1_385-623 | GW_F_ ATGTTAAAGAAACTAGCATGGATAGAAG |
| EDS1_410-623 | GW_F_ ATGTCCTTCAAAGTTTCAAATGAAGAGAATG |
| EDS1_440-623 | GW_F_ ATGAAGAAATGTCAACTTCCAGATGAG |
| EDS1_480-623 | GW_F_ ATGGAAGACACAGGGCCGTAC |
| EDS1_542-623 | GW_F_ ATGTCCGAGTGCGGATCATGC |
| EDS1_542-593 | GW_R_ TGAACCCTCCAGAAATATTTCCTTATC |
| **qPCR** | |
| eEF-1α | F: GATTGCCACACCTCTCACATTGC |
| | R: GGTCCTTCTTGTCCACGCTCTT |
| WRKY18 | F: CGACATACGAAGGGACGCATAAC |
| | R: CCAGATATAGCAGCAGCAAGAGC |
| WRKY40 | F: CTCACTATTGGCGTTACTCGTATGC |
| | R: CCTCTCGGTTATGTTGCTCTTGTT |
| WRKY38 | F: AAGATCAGGATCAGGAGAAGGAAAGC |
| | R: CCAGATGACGATGAAGGAGGATAAGAG |
| WRKY46 | F: GCTTCAACCAAGACAAGAACATCCT |
| | R: TCCAGCAGTGACCATCATCAATAGA |
| WRKY54 | F: GCAAGAGCAAGAGAACAACACCAG |
| | R: GCACAATATGAGTCGAAGAAGGAGGAA |
| WRKY70 | F: ACACCATCTCCGTTCTTGATACCTT |
| | R: TCTGGACTTGCTTTGTTGCCTTG |
| DND1 | F: CTAACCCAGCCCGAACCAGATT |
| | R: CACAACCAACGACAAGCGACTC |
| ERECTA | F: AATACGGTTCTCTTCTTCTTCTCTATCCT |
| | R: CACACCTCTCCAGACACAATAATCC |
| PR1 | F: GCGAGAAGGCTAACTACAACTAC |
| | R: CCTGCATATGATGCTCCTTATTGA |
| PR2 | F: CTTAGCCTCACCACCAATGTTG |
| | R: CGGAATCTGACACCATCTCTGTA |
| PR5 | F: CCACAGACTTCACTCTAAGGAACAAT |
| | R: GTAACCATCTACGAGGCTCACATC |

| | |
|---|---|
| WRKY53 | F: TGACTGGTTCAATCCAACGGTCG |
| | R: GCCGCAAAACTGATGGAAGCATAC |
| WRKY46 | F: GCTTCAACCAAGACAAGAACATCCT |
| | R: TCCAGCAGTGACCATCATCAATAGA |
| SARD1 | F: GGAGAAGTAAGTGTGACGGTGAGAA |
| | R: TGTTGATGTGGCGAGAGGAGAG |
| PAD4 | F: TCAATCTTCTCCGCCGTCATTCC |
| | R: AGTGTCCGTACCTCTGATGTTCCT |
| ICS1 | F: CTAACGAGAACGGAAACGGAAAC |
| | R: TACCACCATAGGCACGAATCAG |
| PBS3 | F: CGTCCTGAATTAGCAGACACTATTGAAG |
| | R: CTCCGAAGAACCGTAAGTTGTTGAA |
| **ChIP-qPCR** | |
| Actin2 | F: GTAACATTGTGCTCAGTGGTGGAA |
| | R: CCTGGACCTGCCTCATCATACT |
| *PR1* amplicon genomic loci relative to TSS | |
| F1 (+336/+606) | F: GCGAGAAGGCTAACTACAACTAC |
| | R: CCTGCATATGATGCTCCTTATTGA |
| F2 (-235/-7) | F: GGCAAAGCTACCGATACGAAAC |
| | R: GGGTTCGTAAACATCGCTTATATAGAG |
| F3 (-574/-300) | F: GCATGAAACACTAAGAAACAAATAATTCTTG |
| | R: TGTATATAGTTGTTTCATGTCATTCAGTTG |
| F4 (-765/-578) | F: ACAAAGTGTATACAATGTCAATCGGTG |
| | R: TGAGTATCTCTATCACTCTTGCCTATG |
| F5 (-2417/-2204) | F: ATGTTTGAGGTTGAGTACGATGGAC |
| | R: TGTATTCTCTTTGCTTCTCCACAAAG |
| *PR2* amplicon genomic loci relative to TSS | |
| R1 (-320/-143) | F: CCATAATTGAACTTGGCTTGTGGGT |
| | R: GAATCGCCCAAACCAATTTGGTT |
| R2 (-401/-295) | F: GAACCATCCATGGCACCTTAAAC |
| | R: GACCCACAAGCCAAGTTCAATTATG |
| R3 (-531/-405) | F: CGAACGTGTTTTACAAGGTTGAGAC |
| | R: GAACCACTGTGAACACAAATCATGC |
| R4 (-706/-576) | F: GCACCTTGTGTTTTACTGAACCAAAT |
| | R: GTCGTTTTCCCCCTTCATGATG |
| *PBS3* amplicon genomic loci relative to ATG code | |
| S1 (-255/-135) | F: CACTTCACACGCTAAGGTCAGT |
| | R: ATGGATTTGGCAGTTCGTGAGT |
| S2 (-561/-383) | F: CCATGAAGCCTTAGATCGATGTCAAAGT |
| | R: CCATGGAAAATGTTTTTCTTTTGTCGGC |
| S3 (-890/-753) | F: CGACAAGATGATACTTGCTGAACAATTC |

| | |
|---|---|
| | R: GCAGTATACTAGTACCATGCAAAATTGTG |
| S4 (-1229/-1062) | F: CTTTCATCACACGACTCACTCAACT |
| | R: TACACAAAGACCAGACCACAACAAG |
| S5 (-1345/-1230) | F: ACCAATCTGGAGAAAAGATACGAGAC |
| | R: TGATGGAGACTAGAAATTACAAACGAAGTTTC |
| S6 (-2182/-2068) | F: ATGGACGCTACTTAAATCGACACG |
| | R: TTCTAGCGAGAAAGGACAAGCCAT |
| S7 (-2341/-2222) | F: GTACATTAAATAATAAGATCCCCTTTGATTTATGG |
| | R: TCGGGCCGTATAAGACATTAC |
| S8 (-2843/-2697) | F: GTCAGAAGATGGAGACTACTTGGAAG |
| | R: TGGAGGGATATTTGCCCAGAAATTC |
| *EDS1* amplicon genomic loci relative to TSS | |
| E1 (+312/+599) | F: AGCTTCTGTGGAAATGGCTGTG |
| | R: ACAGACGCCTTTCGAGCAAG |
| E2 (-167/-39) | F: ACCGACACGTGGAAAGCTAAG |
| | R: TACCCAAAATACCCCATCATGAGACC |
| E3 (-355/-239) | F: AACCGAGAGTGACATCTCAAACC |
| | R: TTTGGTCGGAAAGTTTTCTCTGTGT |
| E4 (-562/-418) | F: CTTGTAGATTATCCGGTTTACTTTCCGG |
| | R: GAATGAGGAGATTTGTTCGCTTGGAG |
| E5 (-622/-609) | F: TTCTAGTCCGGTTCATGTAACTTC |
| | R: ACTAACTACACCTTCTTGCCTGG |
| E6 (-1281/-1184) | F: ACACATGCATGTCCTGATTCTTTCG |
| | R: CCCATCTTTGTCACTCTTGATCAGTCT |
| **Genotyping** | |
| GFP | F: AGGGTGAAGGTGATGCAACATACG |
| | R: TCCAGCAGGACCATGTGATCG |
| eGFP | F: AAGCTGACCCTGAAGTTCATCTGC |
| | R: TCCAGCAGGACCATGTGATCG |
| *eds1-2* | F: GACCCTTTCTAGTTTCCTTGAGCTAAGTC |
| | R: CGTCTGTGATCCATTCTCCAAGC |
| | WT, 1065 bp; *eds1-2*, 222 bp. |
| *npr1-2* | F: GGATGATTTCTACAGCGACGCT |
| (CAPS) | R: GTAACCATAGCTTAATGCAGATGGTG |
| Restriction enzyme | FspI: WT, 270 bp and 139 bp; *npr1-2*, 409 bp. |
| *npr1-3* | F: GGCCGACTATGTGTAGAAATACTAGCG |
| (dCAPS) | R: TGAGACGGTCAGGCTCGAGG |
| Restriction enzyme | HhaI: WT, 319 bp and 27 bp; *npr1-3*, 346 bp. |

**Supplementary Table 2. Recombinant DNA**

| Plasmid name | Source |
|---|---|
| pDEST-GBKT7-NPR1 | This paper |
| pDEST-GBKT7-EDS1 | This paper |
| pDEST-GBKT7-PAD4 | This paper |
| pDEST-GBKT7-PAD4_1-340 | This paper |
| pDEST-GBKT7-SAG101 | This paper |
| pDEST-GBKT7-CDK8 | This paper |
| pDEST-GBKT7-EDS1_Δ480-540 | This paper |
| pDEST-GBKT7-EDS1_Δ358-383 | This paper |
| pDEST-GBKT7-EDS1_Δ310-383 | This paper |
| pDEST-GBKT7-EDS1_1-200 | This paper |
| pDEST-GBKT7-EDS1_1-273 (eds1-2) | This paper |
| pDEST-GBKT7-EDS1_1-300 | This paper |
| pDEST-GBKT7-EDS1_310-424 | This paper |
| pDEST-GBKT7-EDS1_310-383 | This paper |
| pDEST-GBKT7-EDS1_Δ310-350 | This paper |
| pDEST-GBKT7-EDS1_310-330 | This paper |
| pDEST-GBKT7-TGA2 | This paper |
| pDEST-GBKT7-TGA3 | This paper |
| pDEST-GBKT7-EDS1_1-120 | This paper |
| pDEST-GBKT7-EDS1_1-350 | This paper |
| pDEST-GBKT7-EDS1_1-384 | This paper |
| pDEST-GBKT7-EDS1_1-424 | This paper |
| pDEST-GBKT7-EDS1_357-623 | This paper |
| pDEST-GBKT7-EDS1_385-623 | This paper |
| pDEST-GBKT7-EDS1_410-623 | This paper |
| pDEST-GBKT7-EDS1_440-623 | This paper |
| pDEST-GBKT7-EDS1_480-623 | This paper |
| pDEST-GBKT7-EDS1_542-623 | This paper |
| pDEST-GBKT7-EDS1_542-593 | This paper |
| pDEST-GADT7-SAG101 | This paper |
| pDEST-GADT7-PAD4 | This paper |
| pDEST-GADT7-EDS1 | This paper |
| pDEST-GADT7-NPR1 | (ref.[34]) |
| pDEST-GADT7-GUS | (ref.[34]) |
| pDEST-GADT7-EDS1 | This paper |
| pDEST-GADT7-NPR1_1-195 (BTB) | This paper |
| pDEST-GADT7-NPR1_Δ60-150 | This paper |
| pDEST-GADT7-NPR1_1-500 | This paper |
| pDEST-GADT7-NPR1_375-593 | This paper |
| pDEST-GADT7-NPR1ANK | This paper |
| pDEST-GADT7-NPR1C521A | (ref.[34]) |
| pDEST-GADT7-NPR1C529A | (ref.[34]) |

| | |
|---|---|
| pDEST-GADT7-npr1-2 | This paper |
| pDEST-GADT7-nim1-2 | (ref.[34]) |
| pDEST-GADT7-npr1-1 | (ref.[34]) |
| pDEST-GADT7-npr1-5 | (ref.[34]) |
| pDEST15-GST-EDS1 | This paper |
| pDEST15-GST-PAD4 | This paper |
| pDEST15-GST-GUS | This paper |
| pET32a-Trx-$His_6$-NPR1 | (ref.[34]) |
| pMDC43-nVenus-EDS1 | This paper |
| pMDC43-nVenus-NPR1 | This paper |
| pMDC43-cCFP-PAD4 | This paper |
| pMDC43-cCFP-EDS1 | This paper |
| pCAMBIA1300-EDS1-nLUC | This paper |
| pCAMBIA1300-cLUC-NPR1 | This paper |
| pCAMBIA1300-cLUC | (ref.[91]) |
| pCAMBIA1300-nLUC | (ref.[91]) |
| pCAMBIA1300-cLUC-RAR1 | (ref.[91]) |
| pCAMBIA1300-SGT1b-nLUC | (ref.[91]) |
| pEarleyGate302-pNPR1:NPR1-3xFlag | This paper |
| pEarleyGate302-pEDS1:EDS1-3xFlag | This paper |
| pEarleyGate302-pEDS1:EDS1-9xMyc | This paper |
| pEarleyGate203-35S:Myc-GUS | This paper |
| pMDC43-35S:GFP-NPR1 | This paper |
| pMDC83-35S:NPR1-GFP | This paper |
| pMDC43-35S:GFP-NPR3 | This paper |
| pMDC43-35S:GFP-NPR4 | This paper |
| pMDC43-35S:GFP-PBS3 | This paper |
| pK7FWG2-35S:EDS1-eGFP | This paper |
| pK7FWG2-GUS-eGFP | This paper |
| pK7FWG2-35S:EDS1(Δ310-330)-eGFP | This paper |
| pK7FWG2-35S:EDS1(Δ331-350)-eGFP | This paper |
| pMDC43-35S:GFP | This paper |
| pMDC83-35S:TGA2-GFP | This paper |
| pEarleyGate201-35S:HA-CDK8 | This paper |
| pMDC83-35S:EDS1-GFP | This paper |
| pEarleyGate203-35S:Myc-GUS | This paper |
